# Pleiotropy promotes divergence when simulating the development and evolution of two interacting organs

**DOI:** 10.1101/2025.04.11.648404

**Authors:** Pascal Hagolani, Marie Sémon, Guillaume Beslon, Sophie Pantalacci

## Abstract

Pleiotropy, which can occur when a gene affects multiple traits, is a central property of living organisms, influencing their response to mutations and their evolutionary trajectories. Despite many studies and discussions, it remains very difficult to reconcile molecular, developmental and quantitative evolutionary genetics viewpoints on pleiotropy and appreciate how much it puts constraints on genetic evolution, phenotypic evolution and adaptation. Here, we revisit this question by simulating evolution *in silico*. Our model captures multiple levels of integration observed in complex organisms: from genes to development to phenotype to fitness, additionally allowing to remove pleiotropy to directly test its effect. We focus on the pleiotropic interactions between two organs, specifically teeth. Fitness is determined from the functional interaction between these two teeth, which are produced by an *in silico* model of tooth morphogenesis. The developmental parameters are produced by a genome consisting of pleiotropic genes which may be influenced by one tooth-specific transcriptional regulator per tooth. Our simulations with and without pleiotropy confirm several acknowledged consequences of pleiotropy. It reduces genetic and phenotypic exploration, and facilitates the rapid accumulation of adaptive mutations in the modular cis-regulatory regions of pleiotropic genes, which are specific to each tooth. Unexpectedly, pleiotropy promotes fitness improvements, morphological complexity, and the accumulation of genetic divergence. Moreover, in our simulations, mutations in pleiotropic genes contributed significantly to adaptation, and removing pleiotropy did not increase the proportion of adaptive mutations. Thus pleiotropy does not act as a conservative force, but a channeling force promoting genetic divergence.

**Author summary:** The notion of “gene pleiotropy”, like the notion of “gene”, crosses different fields of biology, but views differ on how frequent it is and how much it favors conservation or divergence at the molecular, developmental or phenotypic scale. One reason is the discrepancy between viewpoints at these three scales; another is the difficulty of manipulating pleiotropy experimentally. Here, we performed in *silico* evolution experiments with a computational model focusing on the pleiotropy between two organs while bridging the molecular, developmental, phenotypic and organ function scales. We also compared evolutionary scenarios where the two organs share pleiotropic genes or not. Our results reconcile some of these discrepancies and show that even though non-pleiotropic genes diverge faster than pleiotropic genes, the presence of shared pleiotropic genes accelerates overall genetic divergence.

## Introduction

Pleiotropy is the phenomenon whereby a genetic element (a gene, a mutation) affects multiple traits. It is a central property of the genetic organization of organisms, influencing their response to mutations and their evolutionary trajectories. Fisher’s geometric model, the basis for many studies related to pleiotropy [1–7], predicts that pleiotropy poses such a great disadvantage that adaptation is severely reduced [3]. Since then, many more models have been proposed that moderate this view. These studies showed that pleiotropy does not scale with complexity as in Fisher’s model [8], and that intermediate levels of pleiotropy could be advantageous [9–15]. However, in these models pleiotropy is generally imposed by a predefined mapping from genes to traits, rather than emerging from an explicit developmental process.

Here, we focus on the case of complex organisms, in which pleiotropy is particularly difficult to understand because development and homeostasis integrate diverse traits across multiple organs, and fitness integrates many complex traits [16]. In these organisms, pleiotropy has been defined and studied from multiple viewpoints, e.g., evolutionary and functional genomics, quantitative genetics, or evolutionary developmental genetics, which are often difficult to reconcile [17–21]. Intriguingly, pleiotropy is often considered pervasive in molecular genetics and evolutionary genomics [16], whereas quantification of pleiotropy at the phenotypic level by quantitative genetics suggests it is limited [22]. These different fields also highlight different consequences of pleiotropy on evolution.

In fields focused on molecular evolution (e.g., genomics, evolutionary developmental genetics), it is widely assumed that pleiotropy favors genetic conservation [23–25] while modularity—the organization of the genome into relatively independent functional units—enables adaptation [26]. Both pleiotropy and modularity are observed in the functional organization of the genome. Most gene products are used repeatedly in different organs or at different time points during lifetime; hence most coding sequences are considered pleiotropic, whereas their regulatory sequences are considered more modular, decoupling the functional regulation of gene products. The conservation of coding sequences is positively correlated with the level of gene pleiotropy, which is measured, for example, as a function of the number of organs or tissues in which a gene is expressed [27]. In contrast, cis-regulatory sequences, which regulate pleiotropic genes, evolve more rapidly and are considered to be the main drivers of the phenotypic evolution of complex traits such as morphology [26, 28, 29].

However, this view of pleiotropy as a strong constraint is challenged by fields related to quantitative genetics. First, pleiotropy quantified from genetic crosses appears limited [22]. Second, models and experimental findings emphasize that correlated changes between traits can be advantageous, in which case pleiotropic mutations can facilitate adaptation [9, 10, 12–15, 30, 31]. Selection may shape Genotype–Phenotype (GP) maps so as to balance the constraints of pleiotropy and the advantages of correlated trait changes it provides, and to improve evolvability. However, this would require a high degree of pleiotropic genetic variation masked, for example, by epistatic interactions [32]. Under this hypothesis, it has been argued that pleiotropy could promote genetic divergence by favoring the accumulation of mutations that compensate for each other [32, 33].

Two factors contribute to these discrepancies. First, it is not possible to remove pleiotropy experimentally to test its influence. Second, mutations in pleiotropic genes can have limited or even no pleiotropic effects if these are masked by epistatic interactions with other loci, or more generally by the structure of the GP map [32, 34]. The GP map is notoriously difficult to model without strong genetic simplifications [35, 36]. Yet such simplifications often fail to account for how developmental systems bias phenotypic variation and determine which genetic variation is visible to selection. To circumvent these limitations, we developed an *in silico* evolutionary framework in which pleiotropy can be removed, to study how it influences adaptation and phenotypic and genetic evolution.

This framework is based on an explicit Genotype–Phenotype–Fitness (GPF) map inspired by molecular and developmental rules observed in complex organisms. We use simplified organisms with only two organs — one upper and one lower tooth — that interact to determine fitness. These teeth acquire their shape during development through the same pleiotropic genes (gray genes in Fig 1A), but modular regulation is possible thanks to tooth-specific transcriptional regulators (green and orange regulators for the upper and lower teeth, respectively, in Fig 1A). After tooth development, simulated with an *in silico* model of tooth morphogenesis, fitness is determined by how well tooth shapes fit together (occlusion), measured by their contact surface. As detailed below, this setup allows us to focus on the pleiotropy linking the phenotypes of the two teeth, while selection still operates on the two interacting teeth as a whole. Moreover, mutations in the pleiotropic genes can have either pleiotropic or non-pleiotropic effects depending on the genetic background and the structure of the GP map.

**Fig 1.**
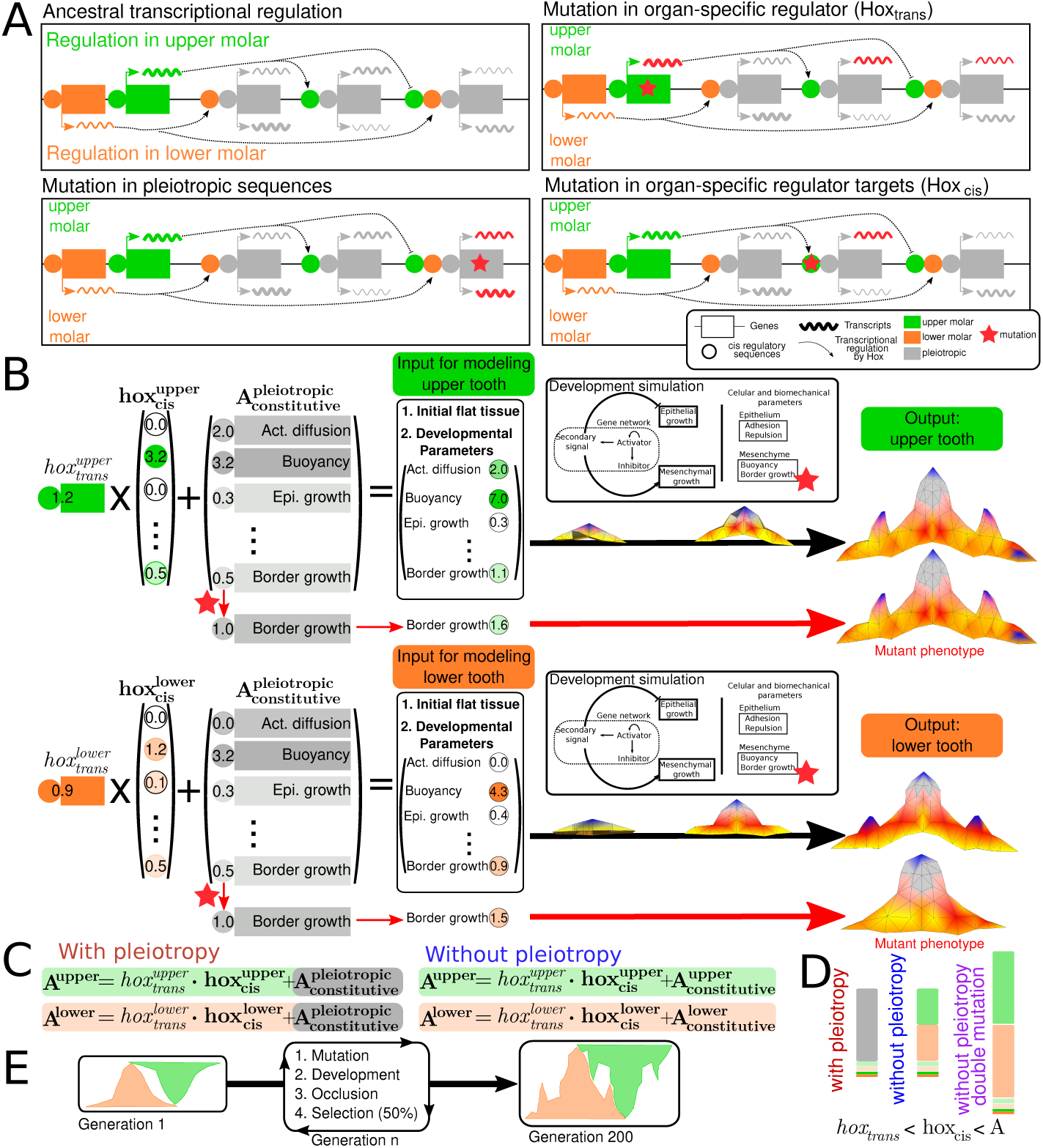
Pleiotropic framework and tooth model. A: Top left: genome modularity shown as transcriptional regulations in the upper and lower teeth. Organ-specific regulators (“Hox” genes in green and orange) bind cis-regulatory regions (green and orange circles) of pleiotropic genes (gray). The other three panels show, respectively, how mutations (red stars) affect three types of genetic elements: pleiotropic (constitutive) sequences, Hox genes and their target cis-regulatory regions. Their effects on transcriptional activity are shown as red transcripts. B: Computational genetic and developmental model. The values of the 21 parameters of the developmental model are computed from the interaction of the Hox and constitutive elements. These parameters then interact to produce the tooth shape through the reciprocal feedback between the epithelium and the mesenchyme. The effect of a pleiotropic mutation on Bgr (boundary growth rate) is shown (red star and arrows). This mutation increases the value of the constitutive element of Bgr. Since Bgr is also regulated by Hox genes in the upper and lower teeth, the developmental model runs with different Bgr values in the two teeth, and in the context of other parameter differences provided by the Hox gene (Epi growth and Buoyancy), resulting in different phenotypic variation for a similar molecular increase. In this example, the mutation has an effect in the lower tooth only, illustrating “hidden” pleiotropy. C: Vectors used to run the developmental model in the with-pleiotropy and without-pleiotropy settings; the without-pleiotropy-double-mutation setting uses the same vectors as without-pleiotropy. D: Proportional mutational influx for the genomic compartments in the three main settings: with-pleiotropy, without-pleiotropy and without-pleiotropy-double-mutation (constitutive, gray; Hox genes, dark color; and cis-regulatory targets, light color. Upper tooth in green and lower in orange). E: Evolutionary model coupled with the developmental model. In each generation, the genomic elements can mutate and the new genomes are passed to the developmental model, which provides the tooth shapes. These shapes are then used/to calculate the occlusion (contact surface between the two teeth); the 50% of individuals with the best occlusion contribute to the next generation.

Our framework has several advantages. First, despite its simplicity, it captures a key property of complex organisms, in which the genetic modularity of body parts is mediated by master transcriptional regulators, such as Hox and Hox-related genes. These genes are specifically expressed in certain regions of the body and introduce body-part-specific regulation of a shared developmental toolkit [37, 38]. For example, we previously showed that during mouse molar morphogenesis, *Pou3f3* is expressed specifically in upper molars, whereas *Nkx2-3* is expressed specifically in lower molars [39], a finding that inspired the tooth-specific regulators in our model.

Second, our framework reproduces the nonlinear, degenerate, and integrated nature of GPF maps, an important ingredient in biological systems. Indeed, tooth shape, as in real organisms, is the product of a complex GP map, since we use a model of tooth morphogenesis that includes both gene–gene and biomechanical interactions, which influence each other during tooth development [40]. This means that our model captures key developmental properties that can influence evolutionary trajectories, such as nonlinearities and fragmentation of the GP map or developmental bias, as shown previously with this type of model [35, 36, 41]. Furthermore, in our framework, fitness is determined by the interaction between two tooth shapes. Hence, fitness emerges from a Phenotype–Fitness (PF) map in which nonlinear effects and degeneracy can occur. This is essential because selection often acts on integrated functions rather than directly on individual traits. In simulations, selection on integrated functions has been shown to favor the emergence of restricted pleiotropy [42].

Third, our framework uses an explicit functional partitioning of the genetic elements into two categories: pleiotropic genes, and organ-specific elements, namely tooth-specific regulators and their target regions in pleiotropic genes. This way, we can independently assess the effect of mutations of these two functional categories. Importantly, mutations that we categorize as pleiotropic at the molecular level could be non-pleiotropic at the phenotypic level, if they either have no effect on the phenotype or affect only one of the two teeth (see example in Fig 1B): this is what we refer to as “hidden” pleiotropy. Indeed, in our model phenotypic effects on tooth shape and possible advantageous/deleterious/neutral effects on fitness are emergent properties of the GPF map. This allows us (1) to compare the rates of evolution of pleiotropic *versus* organ-specific elements and (2) to assess the effect of gene pleiotropy on molecular and phenotypic evolution. Specifically, we can compare evolutionary outcomes of this model, in which teeth share pleiotropic genes possibly regulated by tooth-specific genetic elements, with a control model where each of the two teeth possesses its own set of developmental genes (corresponding to a fully modular setting, with no pleiotropy between the upper and lower teeth).

We found that, although gene pleiotropy reduces phenotypic and genotypic variation, it does not hinder evolution; rather, it promotes it. Indeed, the interplay between pleiotropic and modular parts of the genome channels the exploration of the fitness landscape through a mechanism akin to the path of least resistance [43, 44], resulting in a greater accumulation of adaptive mutations than in the fully modular model.

## Results

### Modeling framework to evaluate how the pleiotropy linking two teeth impacts the evolutionary dynamics

During serial organ development, Hox-like master genes are expressed in specific organs, where they regulate pleiotropic genes through organ-specific cis-regulatory target sequences. The interactions between these genetic elements induce organ-specific differences in gene expression, which affect how cells and tissues interact chemically as well as biophysically over the course of development, resulting in organ-specific morphologies. Our model captures this modular organization for two molars (Fig 1A).

During evolution, mutations may fall in three different genomic compartments, corresponding to these three types of genetic elements: Hox genes, Hox-targeted regulatory sequences and pleiotropic genes (Fig 1A). Accordingly, mutations differ in their molecular effects — being either specific to one tooth or shared by the two teeth (Fig 1B–C) — and in their probability of occurring, which depends on both the per-base mutation rate and the size of the genomic compartments (Fig 1D).

Whatever their nature, mutations change the phenotype only if they change the output of developmental mechanisms. Our evolutionary simulations therefore embed a model of upper and lower tooth development [40], comprising 21 parameters that account for different biological processes (Fig 1B), within a genetic framework designed to disentangle the effects of mutations falling in the three types of genetic elements.

For each tooth and each biological process, we model the proximal molecular Hox-dependent regulation as an additive interaction — the simplest form — between Hox and pleiotropic genetic elements, the latter referred to below as “constitutive” elements. This takes the form of the addition of two vectors: one for the constitutive genetic elements 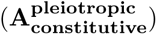 and one for the cis-regulatory genetic elements (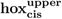 or 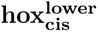). The latter is multiplied by 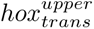 or 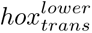, respectively, to model the trans-activity of the Hox regulator (Fig 1C).

The model thus distinguishes three types of mutations (Fig 1A), with different mutational influxes for each type of genetic element (Fig 1D):

- Mutations in the Hox master genes (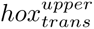 and 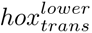), which simultaneously affect the regulation of all target genes, i.e., all target parameters in the model (Fig 1A). Note that elements of 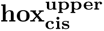 equal to zero are not affected by changes in 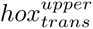. These mutations are the rarest, because in real organisms a single regulator gene is generally involved.
- Mutations in the Hox-cis sequences (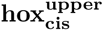 and 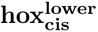), which affect the regulation of specific target genes, and therefore a single parameter in the tooth model (Fig 1A). Because in real organisms Hox genes target several genes contributing to the same biological process, such mutations are more frequent than mutations in the Hox master genes. In our default setting, the mutation rate for each **hox_cis_** element is twice as high as for the *hox_trans_* elements.
- Mutations in the constitutive genetic elements (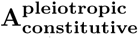), which introduce pleiotropic molecular variation in both teeth (Fig 1A). Note that even though the pleiotropic variation occurs in both teeth simultaneously, the two teeth develop in different epistatic contexts provided by the Hox elements (see the example of the Bgr developmental parameter in Fig 1B). Because in real organisms biological processes are carried out by a large number of pleiotropic genes, such mutations are even more frequent. In our default setting, the mutation rate for the elements is twenty times higher than for the 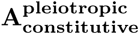 elements.

The genetic elements and mutations described above correspond to the default “with-pleiotropy” setting. To better understand the effect of pleiotropy, we compared this setting to a fully modular model, the “without-pleiotropy” setting (Fig 1C). In the latter, the Hox genes still interact with the constitutive genetic elements to regulate the development of each tooth, but the vector 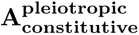 is replaced by two tooth-specific vectors, 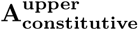 and 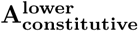 (in green and orange in Fig 1C), such that no genetic element affects both teeth (see below for how mutation rates are adjusted).

In both settings, once the parameter values have been computed from the genetic elements, **A^upper^** and **A^lower^** (Fig 1C) are used to parameterize the tooth developmental model to simulate the development of the upper and lower teeth (Methods, Section Developmental model). We then compute occlusion as the optimal contact surface between the upper and lower teeth (Methods, Section 2D profiles and occlusion); a larger surface indicates better occlusion and thus higher fitness, which determines selection.

At replication, the genome can mutate, each type of compartment having its own rate, so that by the end of a simulation, the total number of mutations is distributed unevenly among the three compartments (Fig 1D; Methods, Section Mutation rates). As the two settings differ in the number of **A_constitutive_** elements (21 for the with-pleiotropy setting versus 42 for the without-pleiotropy setting), using the same mutation rate would result in twice as many constitutive mutations in the latter due to its larger mutational target. To correct for this, we divided the mutation rate of the constitutive elements by two in the without-pleiotropy setting. To assess the effect of this normalization, we also introduced a third setting (“without-pleiotropy-double-mutation” setting), in which the per-element mutation rate is kept identical to that used in the with-pleiotropy setting despite the doubled mutational target.

The model is embedded in an evolutionary loop (Fig 1E; Methods, Section Evolutionary simulations), which we initialize with simple unicuspid teeth that differ between the upper and lower jaws, as seen in the synapsid ancestors of mammals. These initial teeth in our model are already regulated by Hox genes. This is justified because teeth share the Hox-like genes with the upper or lower jaw, which is also an ancestral condition (Methods, Section Initial populations). Each generation comprises 200 individuals, with truncation selection with a 50% threshold. This selection scheme ensures a constant selective pressure, regardless of the distribution of fitness values in the population, while the 50% threshold helps maintain genetic diversity within the population. For each setting — with-pleiotropy, without-pleiotropy, and without-pleiotropy-double-mutation — we ran simulations lasting 200 generations, selecting for improved occlusion. We used 10 independent starting conditions, i.e., 10 different pairs of initial teeth, each with its own configuration of Hox and constitutive elements (Methods, Section Initial populations; SFig 1), and ran 10 replicates for each starting condition, for a total of 100 simulations per setting. In each generation, we recorded fitness and other key phenotypic traits, such as the number of cusps in each tooth. By comparing the final occlusion and the fitness trajectories in the three settings, we analyzed the effects of pleiotropy and modularity on evolution. Since the with-pleiotropy setting contains both types of genetic elements, we also examined the specific contributions of pleiotropic and tooth-specific (modular) genetic elements to evolutionary dynamics.

### Populations simulated with pleiotropy reach better occlusion and greater morphological change

During simulations, occlusion generally increases, with notable changes in tooth morphology and cusp count over time (Fig 2A–B).

**Fig 2.**
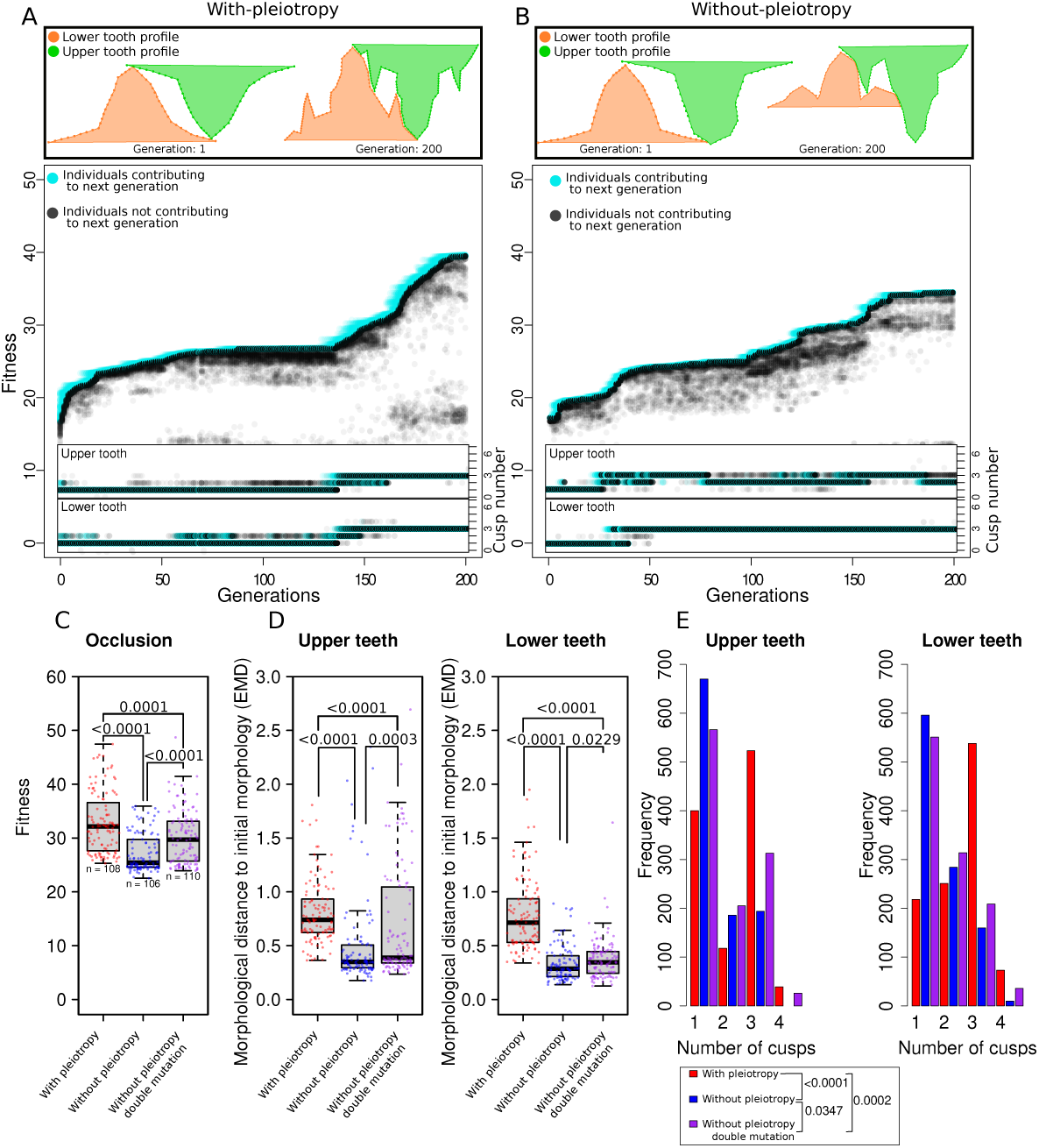
Populations simulated with pleiotropy reach better occlusion values and greater morphological change. A, B: Example simulations in the with-pleiotropy setting (A) and without-pleiotropy setting (B). Top: 2D profiles of the upper and lower teeth in the first (left) and last (right) generations, positioned for optimal occlusion. Note the increase in cusp count and improvement in occlusion. Bottom: Fitness (occlusion) over generations, with the top 50% of individuals (light blue) contributing to the next generation, while the rest (black) do not. The inset shows cusp count changes over generations. C: Mean fitness (occlusion) of the ten best individuals in the last generation for each simulation, per setting. The with-pleiotropy setting achieves significantly better occlusion on average than the without-pleiotropy settings. Average fitness is also higher in the without-pleiotropy-double-mutation setting than in the without-pleiotropy setting. D: Mean morphological distance between the first and last generations of the ten best individuals of each simulation, measured as the Euclidean Morphological Distance (EMD; Methods, Section Euclidean minimal distance (EMD)). The with-pleiotropy setting shows significantly greater morphological change than both without-pleiotropy settings. E: Cusp count per tooth in the last generation. Teeth with more cusps are significantly more prevalent in the with-pleiotropy setting than in either without-pleiotropy setting (upper and lower cusp counts combined for the statistical test). C–E: Pairwise Wilcoxon rank-sum tests (see Methods); *p*-values are shown in the figure.

On average, with-pleiotropy populations achieve significantly higher occlusion than those without pleiotropy, outperforming even the without-pleiotropy-double-mutation setting (Fig 2C). Within each setting, the 10 starting conditions do not significantly affect the outcomes of the simulations (see SFig 2, S1 Table), and the results are robust to different mutation rates — five times lower or higher — and different relative mutation rates between the genetic elements: increasing the mutation rate of Hox elements to the level of constitutive elements (see SFig 3 and SFig 4). Note that, increasing the mutation rate of Hox elements to these extreme levels gives tooth-specific (modular) variation more opportunity to arise (SFig 4), and combined with different initiations of the Hox gene across replicates, tested the robustness of our results to different scenarios of modularity.

**Fig 3.**
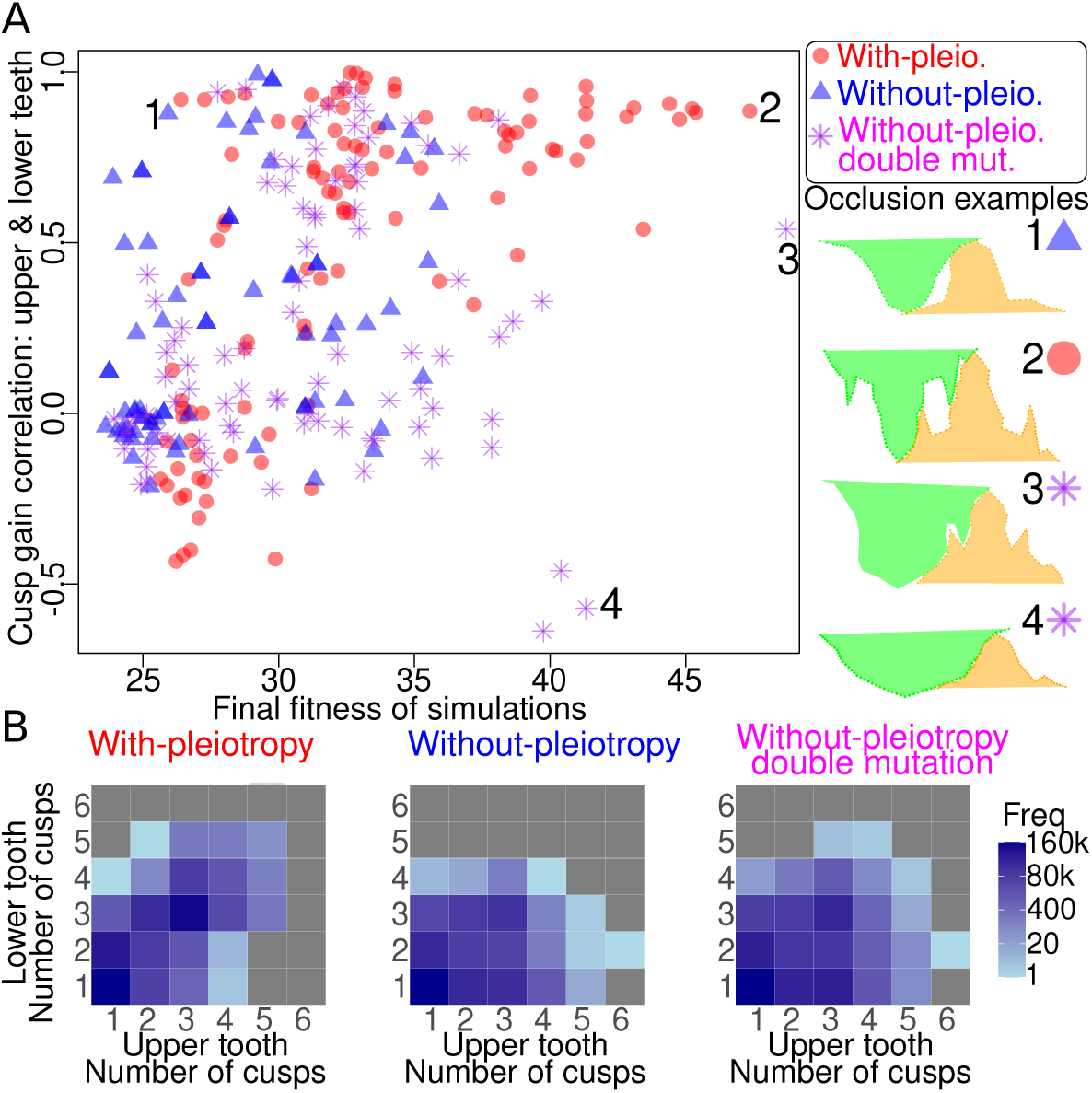
Interaction between cusp gain dynamics and pleiotropy during evolution results in better than average fitness. A: Left panel: Each point corresponds to a single simulation. The x-axis shows the average fitness of the top 10 occluding individuals in the last generation, capturing the evolutionary endpoint. The y-axis shows the correlation of cusp change (gain or loss) between the upper and lower teeth along evolution (measured as the Spearman correlation between the average upper and lower cusp counts of the top 50% occluding individuals in each generation). The correlation between these two quantities is stronger in the with-pleiotropy setting (*r_s_* = 0.6457, *p*-value *<* 0.0001) than in the without-pleiotropy (*r_s_* = 0.3833, *p*-value *<* 0.0001) and the without-pleiotropy-double-mutation (*r_s_* = 0.2317, *p*-value = 0.0282) settings. Right panel: four examples of occlusion at the final generation (numbers correspond to labels in the left panel). Initial conditions do not affect these results, as seen in SFig 5. B: For each of the three settings, the correlation between the number of cusps in the upper and lower teeth, pooling all teeth across all generations. Colors indicate the number of teeth. As for the y-axis in panel A, we use the top 50% occluding individuals of each generation, i.e. the phenotypes that contribute to the next generation.

**Fig 4.**
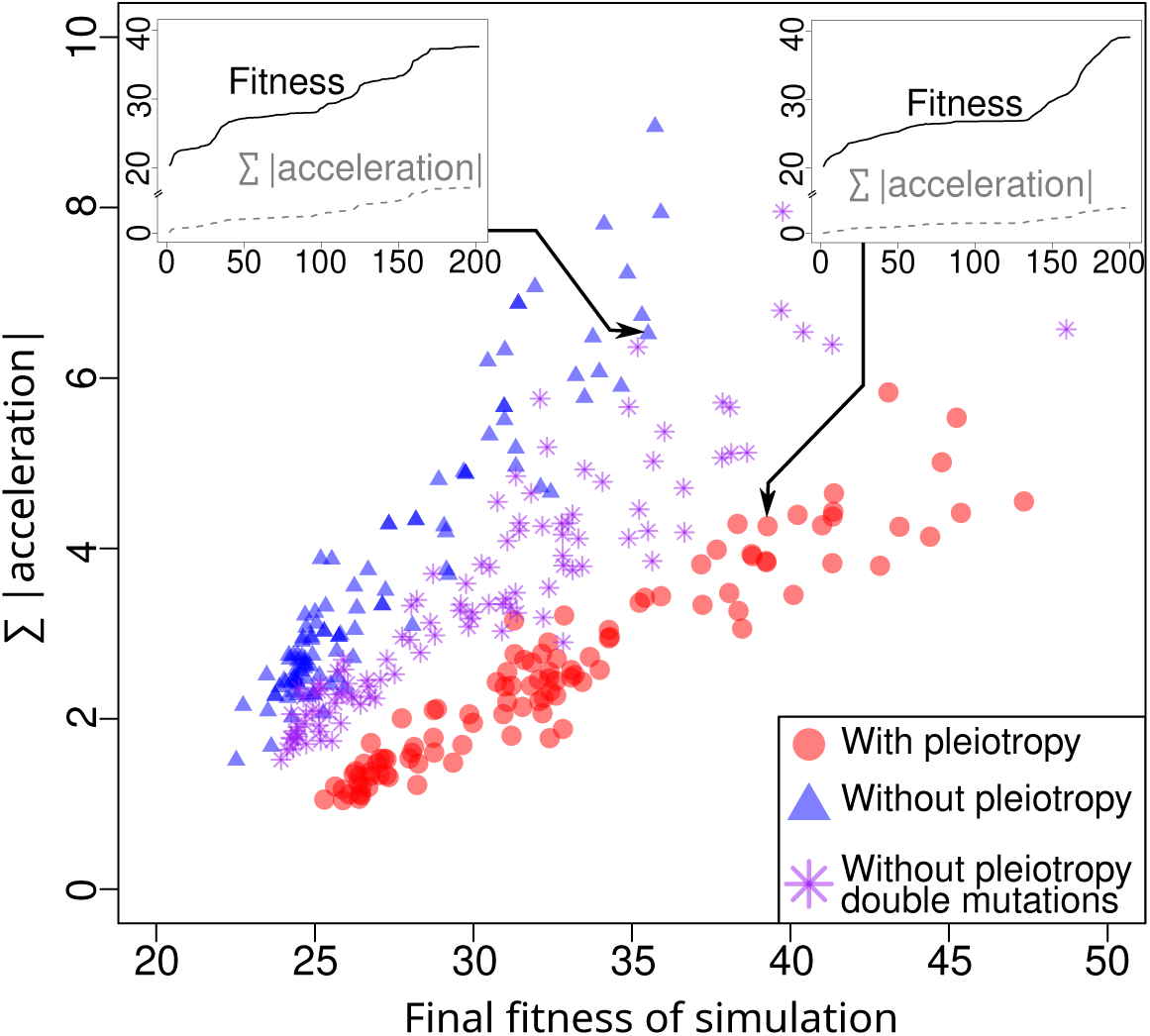
The exploration of the fitness landscape is smoother in the with-pleiotropy setting. We measured, for each simulation of the three settings, the total acceleration in fitness change during evolution (sum of the absolute second derivative of fitness) and plotted it against the final fitness achieved by the simulation. Insets show two examples of fitness trajectories, in the without-pleiotropy setting (left) and in the with-pleiotropy setting (right). In both insets, the black curves show the average fitness of the top 50% occluding individuals at each generation, while the gray curves show the cumulative absolute value of acceleration, computed from the fitness trajectories. For similar final fitness values, simulations in the with-pleiotropy setting show significantly lower total acceleration (ANOVA with final fitness as covariate, *p*-value *<* 0.0001).

Morphology changes more in the with-pleiotropy setting, frequently resulting in multicuspid teeth (Fig 2D–E). The without-pleiotropy settings on average develop fewer cusps in both upper and lower teeth (Fig 2E), often retaining unicuspid teeth similar to the initial generation (Fig 3A, example 1).

To understand why these differences arise, we studied the solutions that lead to better occlusion across settings (SFig 1). Two main solutions emerged: increasing cusp number to expand the contact surface (Fig 3A, example 2) and increasing tooth size (Fig 3A, example 4). A combination of these solutions can optimize occlusion (Fig 3A, example 3). This combined solution, found in the without-pleiotropy-double-mutation setting, achieved the best occlusion observed but occurred in only one simulation.

Because pleiotropy links the evolution of parameters, morphological evolution may be more correlated between upper and lower teeth in the with-pleiotropy setting. Visual inspection of the best final morphologies across all simulations supports this (see SFig 1): in the with-pleiotropy setting, the two teeth of a pair have a similar overall shape, whereas in the without-pleiotropy settings their shapes can differ drastically. This is also true for cusp number. For each simulation, we measured the correlation of cusp change between upper and lower teeth along evolution: this correlation is generally higher in the with-pleiotropy setting than in the without-pleiotropy settings (Fig 3A, y-axis). The same pattern appears when teeth from all generations are pooled (Fig 3B). However, this correlation is far from perfect (see also the inset in Fig 2A). Thus, parallel cusp change in the two teeth occurs more frequently in the with-pleiotropy setting, but not systematically.

We then related this correlation to the final fitness of each simulation (Fig 3A). In all settings, a cluster of simulations combines low cusp-change correlation with poor occlusion, likely corresponding to cases in which tooth morphology remains largely unchanged. Coordinated cusp change is positively correlated with final occlusion in the with-pleiotropy setting (*r_s_* = 0.65), and less so in the without-pleiotropy (*r_s_* = 0.38) and without-pleiotropy-double-mutation (*r_s_* = 0.23) settings (Fig 3A). Starting conditions have no significant effect on these results (SFig 2).

While a strong correlation in cusp change between upper and lower teeth is often advantageous for good occlusion in the with-pleiotropy setting, it is neither sufficient nor necessary. It is not sufficient because, even with pleiotropy, some tooth pairs with strongly correlated changes in cusp number still occlude poorly (e.g., initial condition 3, repetition 1 in SFig 1). Similarly, the teeth in Fig 3A, example 1, show a high correlation in cusp change but remain unicuspid and occlude poorly, because they gained and lost cusps in a correlated way. It is not necessary because, in the two without-pleiotropy settings, other occlusion-enhancing strategies (Fig 3A) do not rely on cusp gain at all. Notably, the best occlusion we observed (Fig 3A, example 3) appeared in the without-pleiotropy-double-mutation setting rather than in the with-pleiotropy setting.

### Fitness increases more steadily in the with-pleiotropy setting

Our previous results show that in our model, pleiotropy does not impede adaptation and that the morphologies of the upper and lower teeth are more correlated in the with-pleiotropy setting than in the two settings evolving without pleiotropy. However, they also show that this correlation does not fully explain the advantage of pleiotropy. To understand this advantage, we analyze how the different settings traverse the fitness landscape in terms of their rate of fitness gain during evolution. To this end, we examined the total acceleration of fitness improvement over time. Fitness acceleration quantifies whether fitness changes are gradual or more saltatory, and thus reveals differences in how the settings evolve. For each evolutionary simulation, we first constructed a fitness trajectory by taking, at each generation, the average fitness of individuals contributing to the subsequent generation, which smooths out individual level variation (black curves in Fig 4 insets). We then computed, for each simulation, the total acceleration as the sum of the absolute second derivative of fitness over its whole trajectory, and plotted it against the final fitness achieved (Fig 4). The three settings clearly separate from one another. For the same final fitness, the with-pleiotropy setting exhibits much lower total acceleration, showing that fitness improves more steadily with pleiotropy than without.

### Parameters in the with-pleiotropy setting change more while exploring a smaller fraction of the parameter space

So far, we compared the different settings based on their fitness and morphological trajectories. To better understand the differences between the with-pleiotropy and without-pleiotropy settings, we analyzed genome evolution by examining changes over time in the different elements of the genome (i.e. the part coded in **A_constitutive_**, which is pleiotropic in the with-pleiotropy setting, and the modular part coded in **hox_cis_**, which includes both 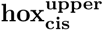 and 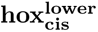.

First, we measured the range between the values at the first and last generations (Δ**_begin_***_−_***_end_**) for each gene in each setting (the raw parameter evolution per initial condition is shown in SFig 6; final fitness, cusp number and cusp-gain correlation do not depend on the initial conditions, Table S1). For the 21 pleiotropic genes, this involves assessing the range in **A_constitutive_** and **hox_cis_** separately. Since different fitness gains could be associated with different levels of gene adaptation, we used a linear model to adjust for fitness differences when comparing ranges in the with-pleiotropy and without-pleiotropy settings (see Methods, Ranges of genetic elements parameter change during evolution). Fig 5A provides an example of the range in the **A_constitutive_** part of the gene that regulates epithelial proliferation. Each point represents a simulation, showing Δ**_begin_***_−_***_end_** for this gene on the *y*-axis (normalized between 0 and 1) and final fitness on the *x*-axis. For each setting, the linear model is shown with trend lines and 95% confidence intervals. Simulations with pleiotropy (in red) consistently show larger ranges than those without (in blue and purple), regardless of fitness. The differences in intercepts between settings estimate the differences in parameter range at equal fitness.

**Fig 5.**
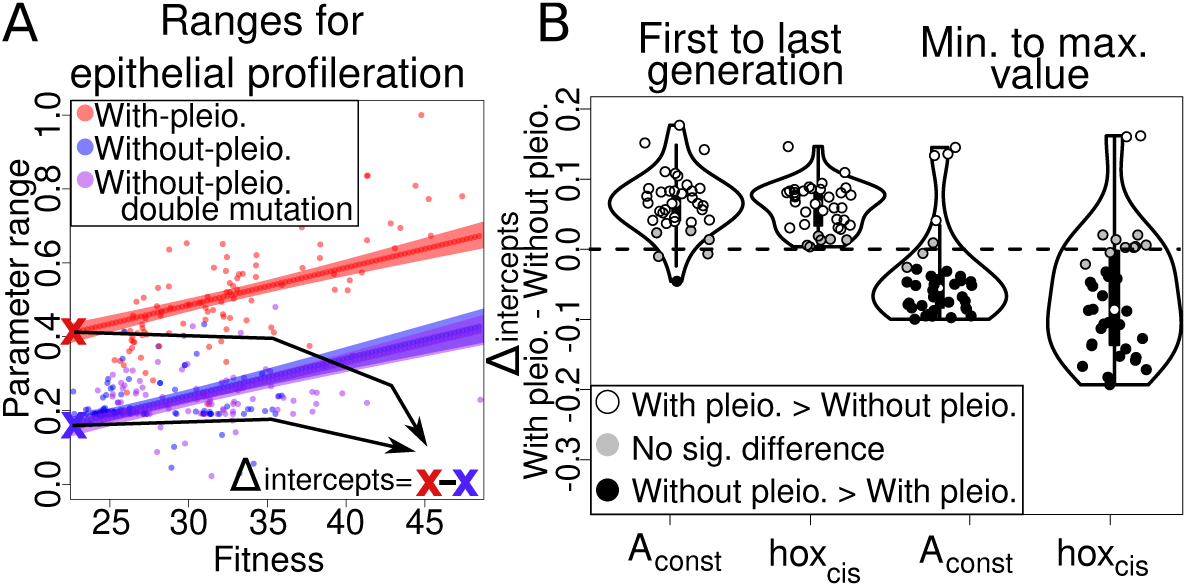
In the with-pleiotropy setting, parameters change more but explore a smaller fraction of the parameter space. For each simulation, we calculated two ranges for the **A_constitutive_** and the upper and lower **hox_cis_** genetic elements: the genetic divergence (Δ_begin_*_−_*_end_; range between the values at the first and last generations) and the overall genetic exploration (Δ_min_*_−_*_max_; range between the lowest and highest values observed across all individuals and generations). A: Example of linear models for Δ_begin_*_−_*_end_ of the **A_constitutive_** element of the gene regulating “Epithelial Proliferation”. Each point represents one evolutionary simulation and colors indicate the setting. The differences between the intercepts of the linear models (crosses) estimate the differences between the settings at equal fitness. B: Comparison of parameter ranges in the with-*versus* without-pleiotropy settings, for Δ_begin_*_−_*_end_ (left violin plots) and Δ_min_*_−_*_max_ (right violin plots), and for the **A_constitutive_** and **hox_cis_** genetic elements. Results for upper and lower teeth are pooled. Each point is the difference between the intercepts of the with- and without-pleiotropy linear models (Δ**_intercepts_**). White circles indicate a significantly larger range in the with-pleiotropy setting, black circles a significantly larger range in the without-pleiotropy setting, and gray circles no significant difference (Tukey’s multiple comparisons of the intercepts). See SFig 9 for the results using the without-pleiotropy-double-mutation setting.

**Fig 6.**
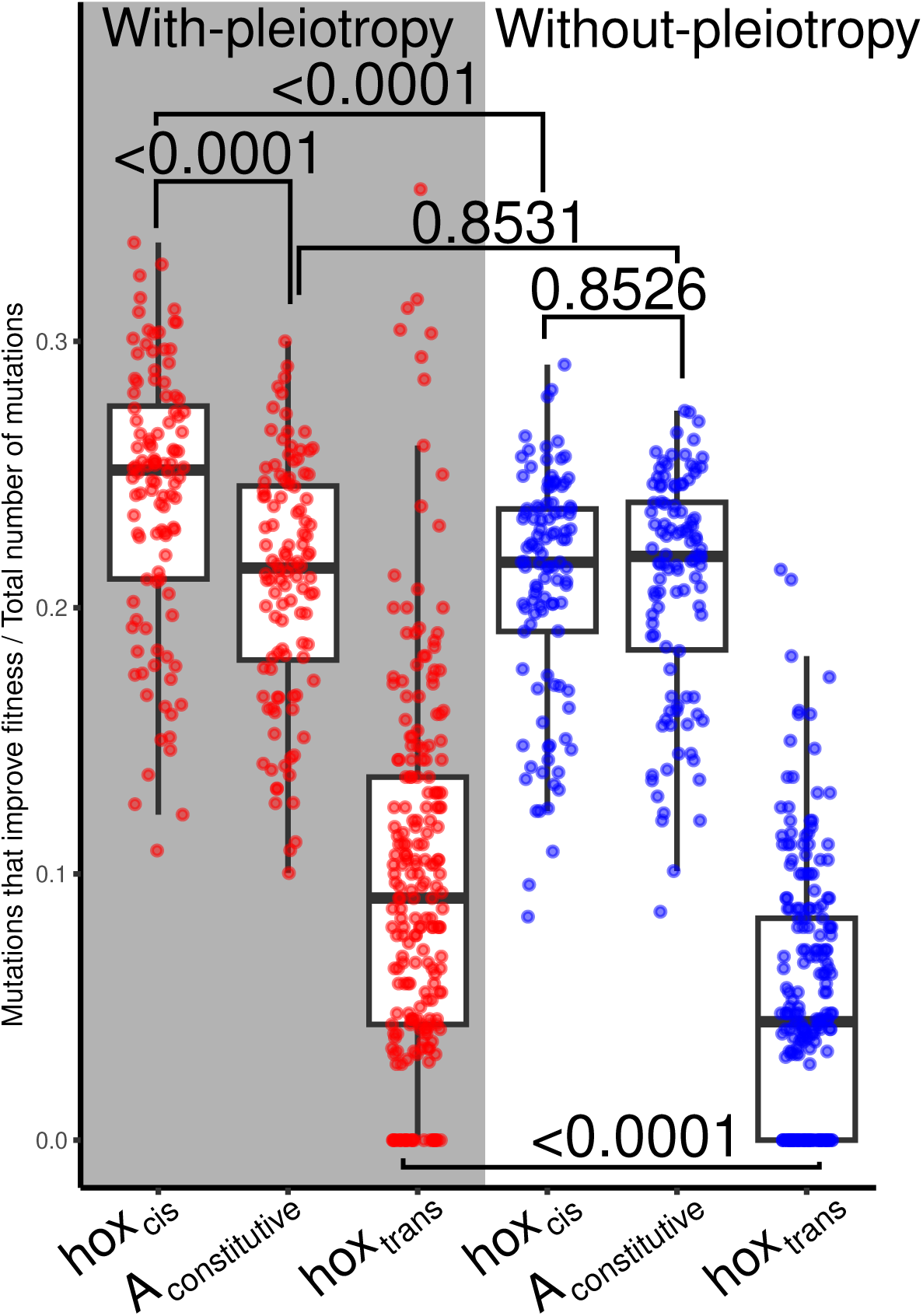
In the with-pleiotropy setting, mutations in the modular parameters (hox_cis_ genetic elements) are more often adaptive, but adaptive mutations also occur in the pleiotropic parameters (A_constitutive_ genetic elements). The y-axis shows, for each simulation, the proportion of mutations that increased fitness and were passed on to the next generation (see Methods, Fraction of adaptive mutations among the different genetic elements). For simplicity, mutations in upper- and lower-specific genetic elements were combined. In the with-pleiotropy setting, the proportion of adaptive mutations in **hox_cis_** genetic elements is significantly higher than in the **A_constitutive_** elements (t-test). In the without-pleiotropy setting, proportions in **hox_cis_** and **A_constitutive_** elements are not significantly different. In the with-pleiotropy setting, the proportion of adaptive mutations in the *hox_trans_* genes is significantly higher than in the without-pleiotropy setting. See SFig 10 for the results including the without-pleiotropy-double-mutation setting.

Fig 5B summarizes the results for the 21 developmental genes, which are shown individually in SFig 7. For most genes, Δ**_begin_***_−_***_end_** (left violin plots) is larger in the with-pleiotropy setting (white circles), suggesting greater accumulation of genetic change during evolution. To test whether this greater accumulation of mutations also involves adaptive mutations, we performed an additional analysis.

Adaptive changes for each gene were estimated from the correlation between the evolution in gene values and improvements in fitness (see Methods, Adaptive change during evolution). The with-pleiotropy setting shows significantly more adaptive changes (see SFig 8). This accumulation of adaptive changes in the with-pleiotropy setting indicates a higher retention of beneficial mutations over time, while the two without-pleiotropy settings show a consistently lower capacity to find or maintain such mutations.

Finally, we calculated the range between each gene’s minimum and maximum values (Δ**_min_***_−_***_max_**) across all individuals and the 200 generations to assess overall parameter space exploration. The right violin plots in Fig 5B show that simulations without pleiotropy generally explore a larger range of gene values, while simulations with pleiotropy exhibit lower Δ**_min_***_−_***_max_** values (black circles), except for 2 genes (white circles). This is consistent with the higher morphological diversity visually observed in the two settings without pleiotropy (SFig 1).

### In the with-pleiotropy setting, single mutations are more often adaptive in modular parameters than in constitutive parameters, but the without-pleiotropy setting reveals that this is not due to a limitation on constitutive parameter adaptive potential

It has been widely argued that morphological adaptation mainly relies on non-pleiotropic mutations that arise in cis-regulatory regions of the genome [45]. To investigate this, we examined the proportion of mutations that increased fitness and were selected and passed on to the next generation. To clearly identify which parameter improved fitness, we focused on mutations affecting one parameter at a time; the proportions are therefore calculated over these single mutations. Fig 6 shows that both the **hox_cis_** and the **A_constitutive_** parts of pleiotropic genes contribute to adaptation. However, in the with-pleiotropy setting (red), mutations in the modular part (**hox_cis_**) contribute significantly more (by 24% relative to **A_constitutive_**) to fitness improvements than mutations in the **A_constitutive_** part. In contrast, in the without-pleiotropy setting (blue) there is no significant difference between the contributions of **hox_cis_** and **A_constitutive_** parts, as expected, since both types of elements play the same role in this setting. Results including the without-pleiotropy-double-mutation setting are shown in SFig 10.

Unexpectedly, the fraction of adaptive mutations in the **A_constitutive_** part did not differ significantly between the with- and without-pleiotropy settings, suggesting that pleiotropy did not limit the adaptive potential of these parameters. Taken together, our results thus suggest that the condition of pleiotropy favors the accumulation of adaptive divergence in modular parameters more than it limits adaptive divergence in pleiotropic parameters.

## Discussion

### A model to bridge pleiotropy across scales and fields of biology

Our model contributes to a long history of simulation studies on pleiotropy and its evolutionary consequences. Our modeling strategy is based on the molecular understanding of pleiotropy and modularity in developmental systems, allowing us to model an explicit GPF map. Because in developmental mechanisms different organizational levels (molecules, cells, tissues) influence each other during development, pleiotropy at the molecular level does not necessarily translate into pleiotropic effects on the phenotype (Fig 1B). Whether the effect of pleiotropic mutations propagates through the physical and temporal scales of development and ultimately affects one or several organs depends on the logic of the developmental mechanism, but also on the genetic background (epistasis). In fact, organ-specific master genes can be viewed as setting up an organ-specific epistatic context for pleiotropic mutations. In our model, we thus introduced pleiotropy explicitly at the molecular level, but it is the organ-specific epistatic interactions determined by the Hox genes and the properties of the GPF map (which exhibits some level of degeneracy at both G-P and P-F levels) that determine whether a mutation is pleiotropic at the phenotypic level. This means that pleiotropy and epistasis emerge at each generation as products of development, as they do in real organisms with complex GP maps [18, 46, 47]. This differs from previous models of pleiotropy, in which levels of pleiotropy and the accompanying epistasis were hard coded (e.g., NK models [48–51]). Even in implementations where pleiotropy and epistasis are specified as independent features, they have been shown to evolve together [50, 52].

Importantly, modularity at the “molecular level” evolves in our model, even though the number of master genes is fixed at two. In the initial conditions, Hox genes regulate only a limited number of parameters, but both the number of controlled parameters and the relative contributions of the **A_constitutive_**, **hox_cis_** and *hox_trans_* elements to each developmental parameter are free to evolve. Consequently, modularity is not fixed but can change over evolutionary time. It may even disappear completely if the contribution of the **hox_cis_** elements evolves to zero, making the teeth completely pleiotropic.

Considering these features, the main strength of our model is that it allows pleiotropy and modularity levels to evolve across explicit physical and temporal scales: from molecules to organs to individuals and populations, throughout developmental and evolutionary time. The trade-off is that we focus only on pleiotropy linking two serial organs. This is undoubtedly a simplification and represents a particular case, whose implications we discuss below.

First, our model includes only two organs, whereas real organisms typically comprise many. Fisher’s model predicts that the cost of pleiotropy increases with the number of traits [3], which might seem to make a two organ model a special case. However, evolutionary simulations have shown that modularity evolves to scale with the number of traits [42], which offsets this cost. In nature, this scaling likely arises because each organ’s development is generally regulated by a specific combination of transcriptional regulators [53], with only some regulators shared between any two organs. Our model compresses this architecture to a single regulator per organ. In other words, both the number of organs and the regulatory complexity that provides modularity are simplified in parallel, so the balance between the cost of pleiotropy and the modularity that offsets it should be broadly preserved.

Second, our model examines serial organs, i.e. organs of similar type. In this context, we found that coordinated evolution can be particularly advantageous. This advantage is likely to be smaller for unrelated organs. Notably, in nature, the extent of shared transcriptional regulation scales with organ relatedness. For example, regulatory regions are almost entirely shared between serial organs but only partially between unrelated organs [27, 54]. Our model, in which the two organs share most of their regulation and differ by only one specific regulator each, therefore corresponds to the case of highly related organs.

In conclusion, despite its simplifying assumptions, our model offers insight into how pleiotropy and modularity evolve wherever interactions between organ shapes determine function and correlated evolution is advantageous. This includes other serial organs, such as limbs, but also potentially unrelated body parts whose shapes must be coordinated. This perspective is timely, as the importance of correlated evolution is increasingly being recognized [55].

### Having pleiotropy can be advantageous, but it channels phenotypic evolution

In our model, a combination of modularity and pleiotropy is on average, advantageous compared to full modularity. Why is it so?

A first possible explanation relates to how we control for the number of mutations (Fig 1). Our without-pleiotropy setting has the same mutational influx in the constitutive part as the with-pleiotropy setting. However, with pleiotropy, a single mutation can potentially affect both teeth, while without pleiotropy, a single mutation affects only one tooth, resulting in fewer opportunities for phenotypic change. To compensate for this effect, we introduced a without-pleiotropy-double-mutation setting in which the mutational influx is doubled, and we also tested other mutation rates. Increasing the mutational influx increases fitness in all settings (SFig 3). However, both the worst and best simulations with pleiotropy still outperform their counterparts in the without-pleiotropy-double-mutation setting. Thus, the better success of the with-pleiotropy setting cannot solely be explained by the increased opportunity for phenotypic change. Changing the same parameter in both teeth at the same time may also matter, as it can facilitate the coordinated evolution of the two teeth.

A second advantage lies in the opportunities for coordinated cusp evolution. Although the path of coordinated cusp evolution is neither necessary nor sufficient for achieving high fitness, and does not lead to the best occlusion between teeth, it is frequently taken by the with-pleiotropy setting, which therefore has a higher proportion of highly successful simulations than the without-pleiotropy settings. Coordinated cusp evolution can also happen in the absence of pleiotropy, but it is much less frequent and requires a high mutation rate, as in the without-pleiotropy-double-mutation setting. Notably, even the least successful simulations of the with-pleiotropy setting, which remain unicuspid, outperform the least successful simulations of the other settings. This suggests an advantage beyond coordinated cusp evolution.

This more general advantage may be reflected in how the with-pleiotropy setting traverses the fitness landscape, with smoother fitness trajectories (Fig 4). Why these trajectories are smoother, and how this helps reach higher fitness, remains unclear and will be studied in the future. However, we noticed that the parameter that differs the most between with-pleiotropy and without-pleiotropy settings, specifically in the **A_constitutive_** part, is “Bgr” (Border Growth rate; SFig 7 and SFig 8). Mutations in this parameter can enlarge the tooth border (Fig 1B), which could easily alter occlusion, even in unicuspid teeth, and may eventually lead to coordinated cusp evolution. We hypothesize that coordinated changes in this parameter through mutations in its **A_constitutive_** part help the with-pleiotropy setting smoothly traverse the fitness landscape.

A further, related point concerns how each setting improves occlusion. In the with-pleiotropy setting, better occlusion is predominantly reached by increasing complexity, that is, by gaining cusps, whereas the without-pleiotropy settings more often achieve it with simpler teeth, for example by increasing tooth size (Fig 3A). This is intriguing because the tooth mutational landscape is shaped by a strong bias toward simple morphologies: most mutations have little or no phenotypic effect, and those that do tend to yield simpler rather than more complex teeth [36, 56]. Despite this simplicity bias, the with-pleiotropy setting is driven into a less accessible region of the morphospace, and does so while exploring a smaller portion of the parameter space (Fig 5B). We suggest that this channeling effect has two origins. First, because additional cusps are not mutationally robust under simplicity bias, they are lost unless actively maintained by selection. Selection, in turn, favors new cusps only if matching cusps appear in the upper and lower teeth within a short evolutionary window, which is precisely what coordinated, pleiotropic change facilitates. Second, developmental models have suggested that complex phenotypes occupy small, relatively isolated patches of the morphospace surrounded by similarly complex forms, whereas simple phenotypes occupy large, well-connected patches [57]. Combined with simplicity bias, this predicts that lineages starting from simple teeth tend to remain simple, with only occasional and relatively abrupt gains in complexity. Once a complex configuration is stabilized by selection, however (as coordinated cusp gain allows in the with-pleiotropy setting), the lineage enters a region of the morphospace where further complexity is more readily accessible, and simpler variants are progressively selected against [58]. Finally, we quantify complexity solely as cusp number, a coarse measure that in our simulations varies mostly between one and three cusps. A finer metric, such as the orientation patch count [59], might capture aspects of complexity that cusp number misses. We did not pursue this here but note it as a possible direction.

In conclusion, the combination of modularity and pleiotropy is advantageous in several ways: it effectively doubles the opportunity for change for a given mutational influx, and helps traverse the fitness landscape more smoothly, presumably by coordinating the evolution of the two teeth and counteracting the simplicity bias. Coordinated cusp evolution is the most successful strategy in the with-pleiotropy setting, whereas more diverse strategies can be found in the without-pleiotropy settings, provided that the mutational influx is sufficient. Thus, while advantageous, the combination of pleiotropy and modularity also biases the range of phenotypic possibilities and channels morphological evolution along fewer paths. This limitation in morphological outcomes is mirrored by a narrower exploration of parameter space with pleiotropy. At this stage, it is unclear whether these limitations are a direct consequence of the genetic coupling imposed by pleiotropy, or an indirect consequence of its benefit: a smoother path to high fitness that does not require large changes. Future work should clarify what drives these differences in evolutionary trajectories.

### Morphological adaptation in modular versus pleiotropic regions

Using our model, we can also quantify where adaptive genetic evolution occurs, distinguishing the pleiotropic and modular parts of the genes. For complex traits such as morphology, it has long been postulated that selection should more frequently act on cis-regulatory sequences than on coding sequences [26]. This prediction rests on two premises. First, morphological change depends on spatiotemporal changes in gene expression during development. Second, because cis-regulatory sequences are thought to be highly modular, changes in these sequences are expected to have far fewer pleiotropic effects than changes in coding sequences.

Consequently, morphological evolution was expected to proceed mainly through non-pleiotropic mutations in highly modular cis-regulatory regions. Two decades of research on the genetics of morphological evolution now paint a somewhat different picture [60].

The predominant role of cis-regulatory mutations in morphological evolution has now been demonstrated in insects, through a compilation of two decades of experimental work [61]. However, case studies typically focus on a single trait and rarely ask whether the cis-regulatory mutation also generated variation in other parts of the organism (but see [14]). Thus, whether morphological evolution occurs predominantly through pleiotropic or non-pleiotropic mutations in these cis-regulatory regions remains an open question, now more than ever, for two reasons. First, work in molecular developmental biology has shown that cis-regulatory sequences are often pleiotropic, a marked shift from earlier views [62]. Second, an increasing number of case studies show that mutations with pleiotropic effects are involved in morphological adaptation, at both micro- and macro-evolutionary scales [9, 14, 15, 63, 64].

Our study adds a quantitative perspective to this question, within the limits of simulating a relatively simple system with only two serial organs (discussed above). In our simulations, changes downstream of the Hox factor (in cis) strongly dominate over changes in the Hox factor itself (in trans). This is consistent with the idea that trans changes are often selected against because of their large effects, but can occasionally be advantageous. Overall, in the with-pleiotropy setting, a larger proportion of mutations is adaptive in the non-pleiotropic parts (**hox_cis_** and *hox_trans_*) than in the pleiotropic part (**A_constitutive_**). Yet, adaptive mutations in the pleiotropic part remain substantial and contribute considerably to genetic evolution. Our study therefore adds to the growing recognition that pleiotropic mutations play an important role in morphological adaptation [9, 11, 14, 15, 65–69].

### Pleiotropy, genetic and developmental divergence

In our simulations, genetic divergence is higher in the with-pleiotropy setting than in the without-pleiotropy settings (Δ**_begin_***_−_***_end_**; Fig 5B and SFig 9), even after correcting for differences in fitness. This may be surprising at first glance, since the dominant view in molecular evolution has regarded pleiotropy as a conservative force that constrains genomic evolution. However, this view largely relies on genome studies comparing pleiotropic and non-pleiotropic regions within a genome, which find greater divergence in non-pleiotropic regions, consistent with the higher proportion of adaptive mutations we find in the non-pleiotropic part (Fig 6). This differs fundamentally from comparing the extent of divergence between lineages evolving with or without pleiotropy, as we do in our simulations.

Pavlicev and Wagner [33] proposed a hypothesis for why pleiotropy could promote genetic divergence: the “selection, pleiotropy, compensation” (SPC) model. According to this model, when selection drives adaptive changes in one organ, it triggers subsequent selection for compensatory mutations that counteract the pleiotropic effects on other organs. Thus, rather than promoting conservation, pleiotropy would facilitate developmental system drift (DSD) and the evolution of the genetic basis of development.

Several lines of evidence in the literature support the idea that pleiotropy promotes divergence. In our previous research on rodent molar adaptation, we found that developmental changes driving shape evolution in one tooth (upper molar) were also present in the development of the other tooth (lower molar), despite its phenotype remaining largely conserved [64]. This showed how pleiotropic changes could be effectively compensated, allowing developmental system drift while maintaining phenotypic stability. Modeling studies of trait evolution under directional and stabilizing selection have also shown rapid developmental system drift when pleiotropic loci responded to selection [70]. Furthermore, analyses of gene expression evolution in mammalian development have revealed that highly pleiotropic genes show as much expression change as less pleiotropic genes [71]. Finally, evidence for pleiotropy-driven compensatory evolution between organs has recently been found in the promoters and enhancers of highly pleiotropic genes, whose sequences evolve faster than those of more organ-specific regulatory regions, even though their activity is maintained over time through compensatory changes in transcription factor binding sites [27].

This work also shows that mutations in the constitutive part of genes are as likely to be adaptive when this part is pleiotropic, in a genome with pleiotropy, as when it is not, in a genome without pleiotropy. Thus, we found no cost of pleiotropy in this comparison. This suggests either that the system has evolved to minimize the burden of pleiotropy (e.g., if very few mutations with a pleiotropic effect at the molecular level have pleiotropic effects at the phenotypic level), or that the cost exists but is efficiently balanced by compensatory mutations, as in the SPC model. Future studies using a framework that allows individual mutations to be traced will be crucial to test these hypotheses and to better understand how the combination of pleiotropy with some modularity channels phenotypic evolution and promotes genetic divergence.

## Materials and methods

### Developmental model

The tooth developmental model we use simulates tooth morphogenesis [40]. It uses a set of parameters and partial differential equations to describe cell behaviors and gene dynamics, and outputs a 3D tooth.

The model integrates gene regulatory dynamics and tissue mechanics [40]. It starts with a flat layer of epithelium overlying the mesenchyme, mimicking the initial stage of tooth development. In the model, cells produce signaling molecules that diffuse throughout the tissue. These signaling molecules regulate various cell behaviors, such as cell division and differentiation. They can also regulate the expression of other gene products. The model includes three types of signaling molecules. The first is an activator, which promotes both its own synthesis and the differentiation of epithelial cells into enamel knots, the key signaling centers in tooth development [72]. Enamel knot cells stop dividing and produce the two other signaling molecules. The second is an inhibitor, which inhibits the activator and prevents the formation of enamel knots too close to each other. The distance between enamel knots therefore depends on the diffusion and decay of both the activator and the inhibitor. Finally, the third is a secondary signal, which is promoted by the activator. This signal promotes differentiation, inhibits epithelial growth, and activates mesenchymal growth. The model also includes growth biases along the antero-posterior and bucco-lingual axes, which can strongly influence tooth morphology. In total, the model includes 21 parameters describing the production, decay, diffusion, and interaction of the different signaling molecules, as well as various biomechanical properties of tooth development, such as internal turgescence pressures on the epithelium or the adhesion between cells. The same developmental model is used for both teeth; it produces an upper tooth by default, and the lower tooth is obtained by multiplying the z-coordinates of all nodes by −1.

### Evolutionary simulations

Each simulation follows a population of 200 individuals over discrete, non-overlapping generations. Each individual carries a genotype composed of **A_constitutive_**, **hox_cis_** and *hox_trans_* elements (see Mutation rates), from which the developmental parameters of each tooth are computed as

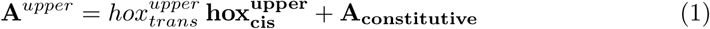

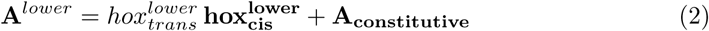

where **A***^upper^*, **A***^lower^*, **hox_cis_** and **A_constitutive_** are vectors of 21 values, one per developmental parameter, and 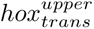 and 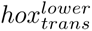 are scalars (Fig 1C). In the with-pleiotropy setting, **A_constitutive_** is shared by both teeth; in the without-pleiotropy settings, it is replaced by two tooth-specific vectors, 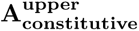 and 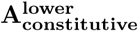.

Each generation consists of four steps. (1) Development: the developmental model is run for each individual, producing an upper and a lower tooth (see Developmental model). (2) Fitness evaluation: during the stabilizing phase, fitness is 1*/*(1 + EMD), where EMD is the Euclidean Morphological Distance (see Euclidean minimal distance (EMD)) between an individual’s teeth and the initial unicuspid teeth, summed over both teeth, so that individuals closer to the initial morphology have higher fitness; during the directional phase, fitness is the occlusion between the upper and lower teeth (see 2D profiles and occlusion). Individuals for which the developmental model fails to produce a final phenotype are assigned a fitness of zero. (3) Selection: the 50% of individuals with the highest fitness (truncation selection) become the parents of the next generation. (4) Reproduction and mutation: reproduction is asexual, and each parent produces two offspring, so that population size remains constant. Each element of an offspring’s genotype mutates independently with the probability given by its mutation rate, and with a mutation size drawn as described in Mutation magnitude. Parents do not survive into the next generation. Each simulation runs for 50 generations of stabilizing selection followed by 200 generations of directional selection (see Initial populations).

### Mutation rates

To modulate pleiotropy, the 42 developmental parameters of the two teeth (21 for each tooth) are encoded in a genetic representation that includes pleiotropic and non-pleiotropic elements. This representation consists of three types of elements: *hox_trans_*, **hox_cis_** and **A_constitutive_**. Biologically, these elements correspond to genetic objects with very different mutational target sizes, resulting in different evolutionary rates. To account for these differences, we assigned a different mutation rate to each type of element. **A_constitutive_** elements represent a large number of genes and gene networks; they summarize a broad range of developmental dynamics into a single parameter. We therefore assigned them the highest mutation rate. **hox_cis_** elements represent cis-regulatory regions, such as enhancers, of specific genes. They correspond to smaller genomic regions so we assigned them a lower mutation rate. *hox_trans_* elements represent single genes, and following the same logic, we assigned them the lowest mutation rate. In the with-pleiotropy setting, the mutation rates are **A_constitutive_**: *µ_p_* = 0.025; **hox_cis_**: *µ_m_* = 0.0025; *hox_trans_*: *µ_t_* = 0.00125. Each rate is the probability that a given element mutates per generation, since mutations occur independently for each element, multiple mutations can occur simultaneously within the same individual.

The without-pleiotropy setting contains twice as many **A_constitutive_** elements, as it includes 2 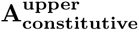 and 21 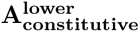 elements. To maintain the same overall mutational pressure as in the with-pleiotropy setting, we halved their mutation rate (*µ_p_*= 0.0125), while the rates of the **hox_cis_** and *hox_trans_* elements remain unchanged (*µ_m_* = 0.0025; *µ_t_* = 0.00125). Finally, to assess the effect of this normalization, the without-pleiotropy-double-mutation setting uses the same mutation rates as the with-pleiotropy setting (*µ_p_* = 0.025; *µ_m_* = 0.0025; *µ_t_* = 0.00125). Because the **A_constitutive_** elements are duplicated, this doubles the number of mutations in these elements per generation. Note that, from the perspective of each tooth rather than of the whole individual, each tooth will then receive roughly the same number of mutations as it does in the with-pleiotropy setting.

### Mutation magnitude

Mutation affects each parameter by a factor *s_i_ ·* (*N* (0, 1)), where *s_i_* is a parameter-specific value for parameter *i* and *N* (0, 1) is a random value drawn from a standard normal distribution. To determine the *s_i_* values, we followed two steps. First, we ensured that the mutational steps were large enough for each parameter to be able to increase tenfold from its initial value within 200 generations. This ensured that all parameters could efficiently explore the parameter space. Second, we verified that, at least around the initial parameter values, the average mutational step had a small but detectable effect on the tooth morphology. Due to the complex interactions intrinsic to the developmental model, this can only be assessed for a limited number of parameter combinations. We therefore focused on the initial parameter values used in the simulations, and measured the Euclidean Morphological Distance (EMD) between the original tooth and the mutated tooth. An EMD = 0 indicates that the morphology did not change. If the average EMD was 0, *s_i_* was increased; if it exceeded 0.2, indicating excessive morphological change, *s_i_* was reduced. This ensured that, at least during the initial stages of evolution, mutations generated sufficient morphological variation to allow evolution, while avoiding excessively large variation that would be likely deleterious.

### Initial populations

We created ten different initial individuals in order to start evolution from different points in the parameter space. Each individual was created by drawing random developmental parameter values, separately for the upper and the lower tooth, until both teeth were unicuspid. Each individual therefore has upper and lower teeth with different parameter values, although both are unicuspid. These values were then split into pleiotropic and modular elements. Each developmental parameter was assigned to be initially regulated by either the upper or the lower modular element. For a parameter regulated by the upper element, **A_constitutive_** takes the value of the lower tooth, 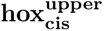 takes the difference between the upper and lower values, and 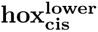 is set to 0; conversely for parameters regulated by the lower element. To choose which parameters are initially regulated by which modular element, we took inspiration from a natural system, reasoning that this could be useful because modularity must emerge from scratch in our model, whereas in nature it is the product of long-term evolution. In mouse molars, the regulatory protein specific to the upper tooth (Pou3f3) is expressed in the mesenchyme, whereas the one specific to the lower tooth (Nkx2-3) is expressed in the epithelium [39]. We therefore assigned the six parameters specific to epithelial or mesenchymal dynamics according to this pattern: in the upper tooth, the modular elements initially regulated the parameters related to mesenchymal dynamics (mesenchymal growth, buoyancy and border growth), and in the lower tooth, those related to epithelial dynamics (epithelial growth, adhesion and repulsion of epithelial cells). The remaining 15 parameters were randomly assigned to either the upper or the lower modular element. This corresponds to a simplified representation of mouse molar development.

Each of the ten initial individuals defines one initial condition. For each, we created a clonal population of 200 individuals (the initial population) and applied 50 generations of stabilizing selection to maintain the unicuspid morphology of both teeth; this stabilizing phase was run once per initial condition. Each resulting population was then used as the starting point for 10 replicate simulations of 200 generations of directional selection to improve occlusion, for each of the three settings. This gives a total of 10 *×* 10 *×* 3 = 300 evolutionary simulations.

### Euclidean minimal distance (EMD)

This measure allows to compare morphologies made of different numbers of cells without requiring an arbitrary selection of landmarks or special morphological features [36]. This is particularly convenient for our study as teeth can have different number of nodes. EMD is defined as the mean distance from each cell in one morphology to the closest cell in the other morphology. This is calculated as follows:

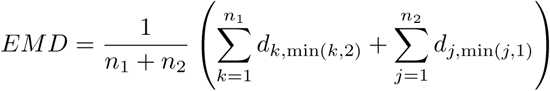

Where *n*_1_ and *n*_2_ are the numbers of nodes in teeth 1 and 2 respectively. *d_k,_*_min(*k,*2)_ is the distance between cell k in tooth 1 and its closest node in tooth 2. *d_j,_*_min(*j,*1)_ is the distance between node j in tooth 2 and its closest node in tooth 1.

### 2D profiles and occlusion

To compute the fitness of an individual, we calculate the contact surface between its upper and lower teeth. This surface defines the occlusion value, the higher the occlusion, the higher the fitness of the individual. Although occlusion is more complex in real organisms, this measurement provides a good proxy for how two artificial teeth evolve to improve their functional interaction. The teeth that we model have 3D shapes, but the main axis of variation is antero-posterior, so they can be accurately represented in two dimensions. To this aim, we first find the extreme cells in the antero-posterior dimension (x-axis). Then we find the tooth cusps by locating the enamel knot cells. In most cases, each cusp only has one enamel knot cell. However, occasionally several enamel knot cells can be associated with the same cusp. To filter-out these cases, the Cartesian coordinates of enamel knot cells that are close to each other (*d <* 1) are averaged and counted as a single enamel knot. Next, we construct the 2D profile, starting from the cusp tips and bounded by the extreme antero-posterior points. We handle each cusp one by one. For each cusp, we trace one flank downhill along the antero-posterior (x) axis: from the cusp tip, we repeatedly step to the tallest neighboring cell with a lower x-value, stopping when no such neighbor remains or we reach the minimum x. Requiring x to decrease at each step keeps the path descending and prevents backtracking. We trace the opposite flank the same way, stepping instead to cells with increasing x. Combining all of the recorded coordinates of the selected cells and dropping the y-coordinate results in a 2D profile of the tooth, which summarizes its space occupancy along the antero-posterior axis. Using the 2D profiles of the two teeth, we calculate the contact surface between them.

To obtain facing upper and lower teeth, the z-coordinates of the lower tooth are multiplied by −1 (see Developmental model), which keeps the antero-posterior and bucco-lingual biases the same in both teeth. For a detailed explanation of how the occlusion is calculated, see SFig 11.

### Ranges of genetic elements parameter change during evolution

For each simulation and each genetic element (**A_constitutive_**, 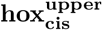 and 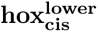, we computed two ranges of parameter values. The genetic divergence, Δ**_begin_***_−_***_end_**, is the absolute difference between the average parameter values of the individuals in the population at the first and the last generation of directional selection. The overall genetic exploration, Δ**_min_***_−_***_max_**, is the difference between the highest and lowest values observed across all individuals and all generations. To make parameters with different scales comparable, each parameter was min-max normalized to the interval [0, 1], jointly across the with- and without-pleiotropy settings, so that the same normalization applies to all settings.

Because simulations that reach higher fitness may also accumulate more parameter change, we compared ranges between settings while accounting for final fitness. For each parameter and genetic element, we fitted a linear model with the range as response, and final fitness and setting as explanatory variables.

The difference in intercepts between two settings, Δ**_intercepts_**, estimates the difference in parameter range at equal fitness (Fig 5A). Positive values of Δ**_intercepts_** indicate a larger range in the with-pleiotropy setting.

Differences between settings were tested with Tukey’s all-pairwise comparisons of the setting effects in the same model, which adjusts for the three pairwise comparisons within each parameter. Upper- and lower-tooth results are shown pooled in Fig 5B. Fig 5 compares the with- and without-pleiotropy settings; the comparison with the without-pleiotropy-double-mutation setting is shown in SFig 9.

### Adaptive change during evolution

We used two complementary measures of adaptation. The measure described here identifies parameters whose evolution is associated with fitness improvement over evolutionary time (SFig 8); the second, described in Fraction of adaptive mutations among the different genetic elements, quantifies the proportion of single mutations that improve fitness (Fig 6).

Fitness changes may result from the simultaneous variation of several parameters, some parameters can hitchhike with others that improve fitness, and several parameters may need to change together to improve fitness. Individual parameter changes therefore cannot be directly attributed to fitness gains. Instead, we measured how consistently each parameter’s evolution is associated with fitness improvement. For each simulation, we computed at each generation the mean parameter value and the mean fitness of the top 50% occluding individuals. We then computed the Pearson correlation between these two time series in rolling windows of 50 generations, shifted by one generation, and classified a window as adaptive for a given parameter if the correlation exceeded r=0.7. The proportion of adaptive windows per simulation measures how consistently a parameter’s evolution tracks fitness improvement, which reduces the influence of hitchhiking and stochastic dynamics at any single time point.

To compare settings while accounting for fitness, we fitted, for each parameter, a generalized linear model with a binomial distribution, with the number of adaptive windows out of the total number of windows per simulation as response, and setting and final fitness (mean fitness of the ten best individuals) as explanatory variables. The predicted proportions of adaptive windows at equal fitness from this model were used to estimate differences between settings. The significance of these differences was tested with Tukey’s all-pairwise comparisons in a linear model of the number of adaptive windows, with setting and final fitness as explanatory variables.

### Fraction of adaptive mutations among the different genetic elements

For each type of genetic element (**A_constitutive_**, **hox_cis_** and *hox_trans_*), we computed the fraction of adaptive mutations as the number of mutations that improved fitness and were passed on to the next generation, divided by the total number of mutations. Mutations that disrupted development (i.e., for which the model failed to produce a final phenotype) were included in the total number of mutations. For simplicity, we only considered mutation events that changed a single parameter, so that any change in fitness can be attributed to that parameter. Upper- and lower-specific elements of the same type were combined. Differences between genetic elements and between settings were tested with t-tests.

### Statistical tests

Statistical analyses were performed in R version 4.6.1 [73], using the packages car, coin, dunn.test, multcomp and visreg. Normality and homoscedasticity were checked with Shapiro–Wilk and Levene’s tests, respectively (Table S3). Parametric tests were used when their assumptions were met, or when models with several explanatory variables (e.g., fitness as a covariate) were required; otherwise, non-parametric tests were used. The names of the tests are provided in the main text and figure captions. *p*-values are generally displayed in the figures, and exact *p*-values and full statistical results are provided in Table S4. All analyses and figures were produced with custom R scripts. Data available at https://doi.org/10.5281/zenodo.23952313 and code at https://github.com/hagolani/PleiotropyDivergenceTwoOrgans and https://doi.org/10.5281/zenodo.23954307.

## Acknowledgments

This work was supported by a grant from IDEX Lyon ANR-16-IDEX-0005, and an European Council Research grant (ERC 2022 COG PLEIOTROPY 101088398). Salaries were supported by the Centre National de la Recherche Scientifique, the Ecole Normale Supérieure de Lyon and the INSA de Lyon. HPC resources were provided by the CBPsmn (PSMN, Pôle Scientifique de Modélisation Numérique) of the ENS de Lyon. We also thank Lisandro Milocco, Benjamin Audit, Sabrina Renaud, Teemu Häkkinen and the members of the BioTiC team for their considerable advice and valuable comments. Finally, we thank reviewers for their detailed comments which helped to improve this manuscript.

## Author contributions

Conceptualization MS SP Formal Analysis MS PH Funding Acquisition MS SP Investigation PH GB MS SP Methodology PH GB MS SP Vizualisation PH GB Software PH Supervision MS SP Writting (Draft) PH MS SP Writting (Review/Editing) PH GB MS SP

## Financial Disclosure

This work was supported by an European Research Council grant (ERC 2022 COG PLEIOTROPY 101088398 to MS; PH) and an IDEX Lyon grant (ANR-16-IDEX-0005 to MS). Salaries were supported by the Centre National de la Recherche Scientifique, the Ecole Normale Supérieure de Lyon and the INSA de Lyon. The funders had no role in study design, data collection and analysis, decision to publish, or preparation of the manuscript.

## Supporting information

**Supplementary figure 1.**
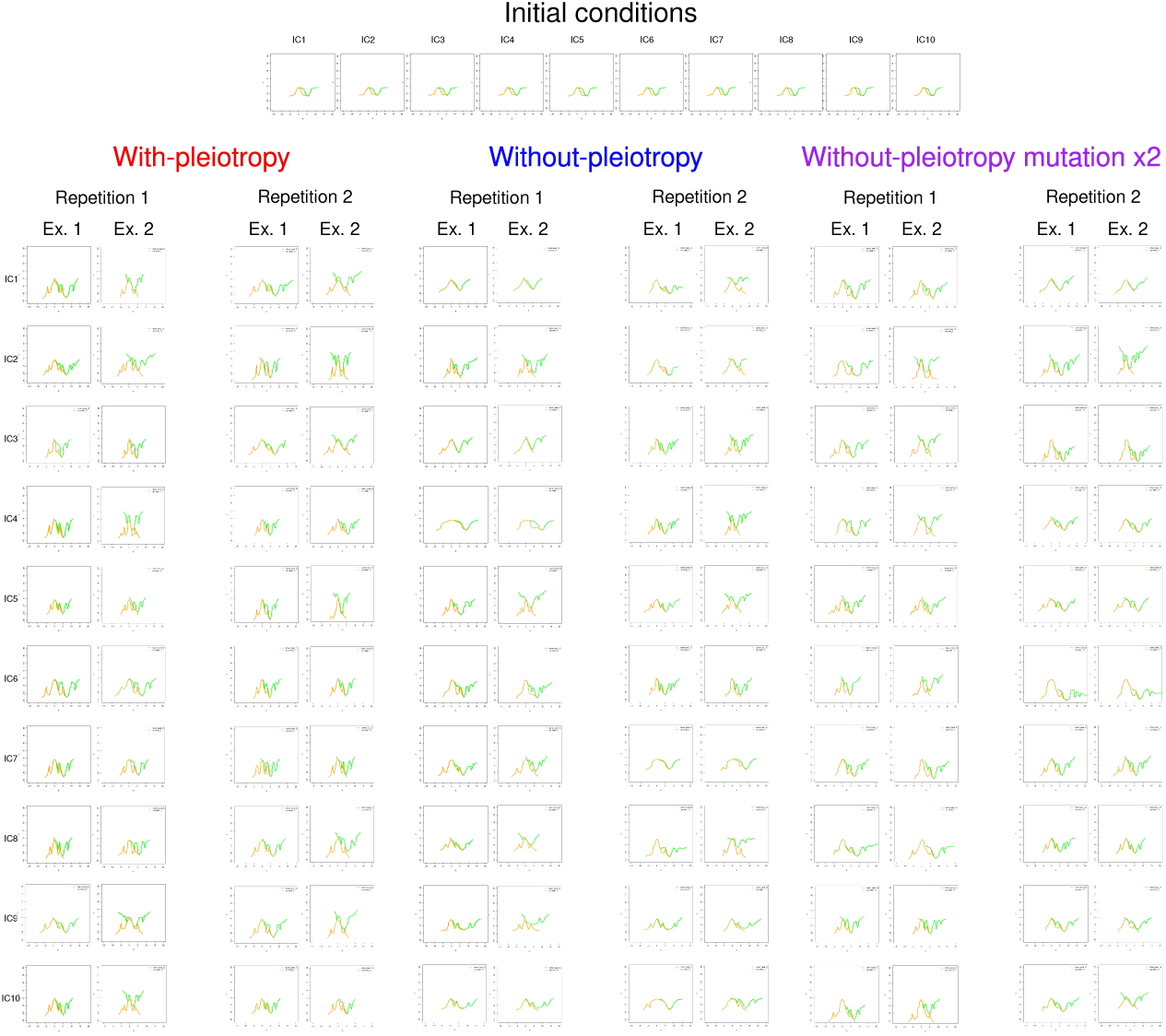
Zoo of the 10 initial conditions and examples of final morphologies. The top row of morphologies shows the 10 different initial conditions, each used to start 10 independent simulations per setting. All teeth are initially unicuspid, but the 10 pairs differ slightly in morphology, and within each pair the upper and lower teeth also differ slightly. Below, for each initial condition, we show morphologies from the last generation of the corresponding simulations. For each setting, we show two randomly chosen simulations out of the ten replicates (labeled repetition 1 and 2). For each repetition, we show two individuals from the final population: one with high occlusion (Ex.1, left) and one with lower occlusion (Ex.2, right). In the with-pleiotropy setting, upper and lower teeth tend to have a similar overall shape, with minor differences, whereas in the without-pleiotropy settings, the upper and lower teeth of an individual frequently differ markedly.

**Supplementary figure 2.**
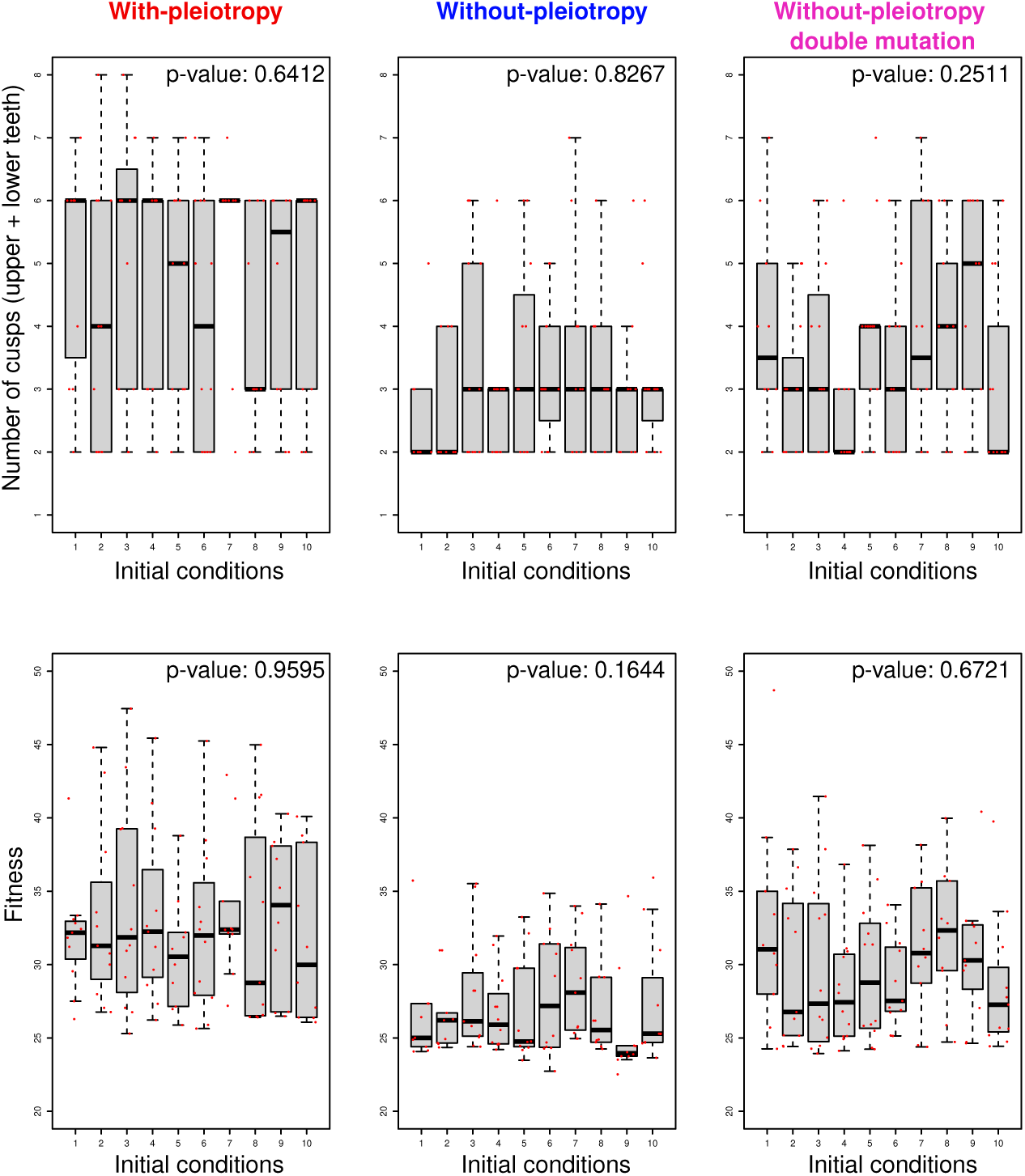
Effect of initial conditions. For each of the 10 initial conditions, we plot the final fitness and the combined number of cusps of the upper and lower teeth in each of the 10 replicate simulations. Neither cusp number nor final fitness differed significantly between initial conditions (Kruskal–Wallis tests; *p*-values shown on each panel and in Table S1 Table).

**Supplementary figure 3.**
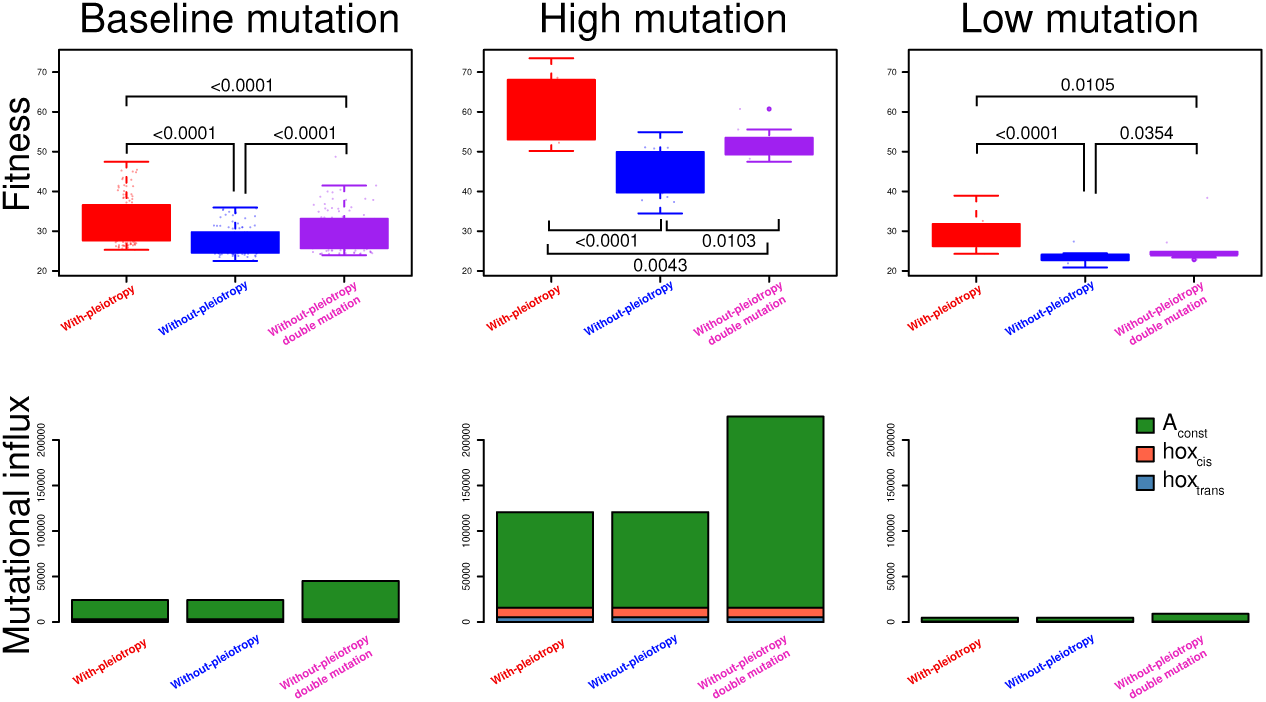
Mutational influx. The with-pleiotropy setting reaches higher fitness than the without-pleiotropy settings across different mutation rates. In addition to the baseline rates, we tested a high mutation rate, five times the baseline, and a low mutation rate, one fifth of the baseline. The resulting mutational influx is shown in the bottom row. The fitness advantage of the with-pleiotropy setting over both without-pleiotropy settings remains significant at all mutation rates (Wilcoxon rank-sum tests; *p*-values are shown in the figure).

**Supplementary figure 4.**
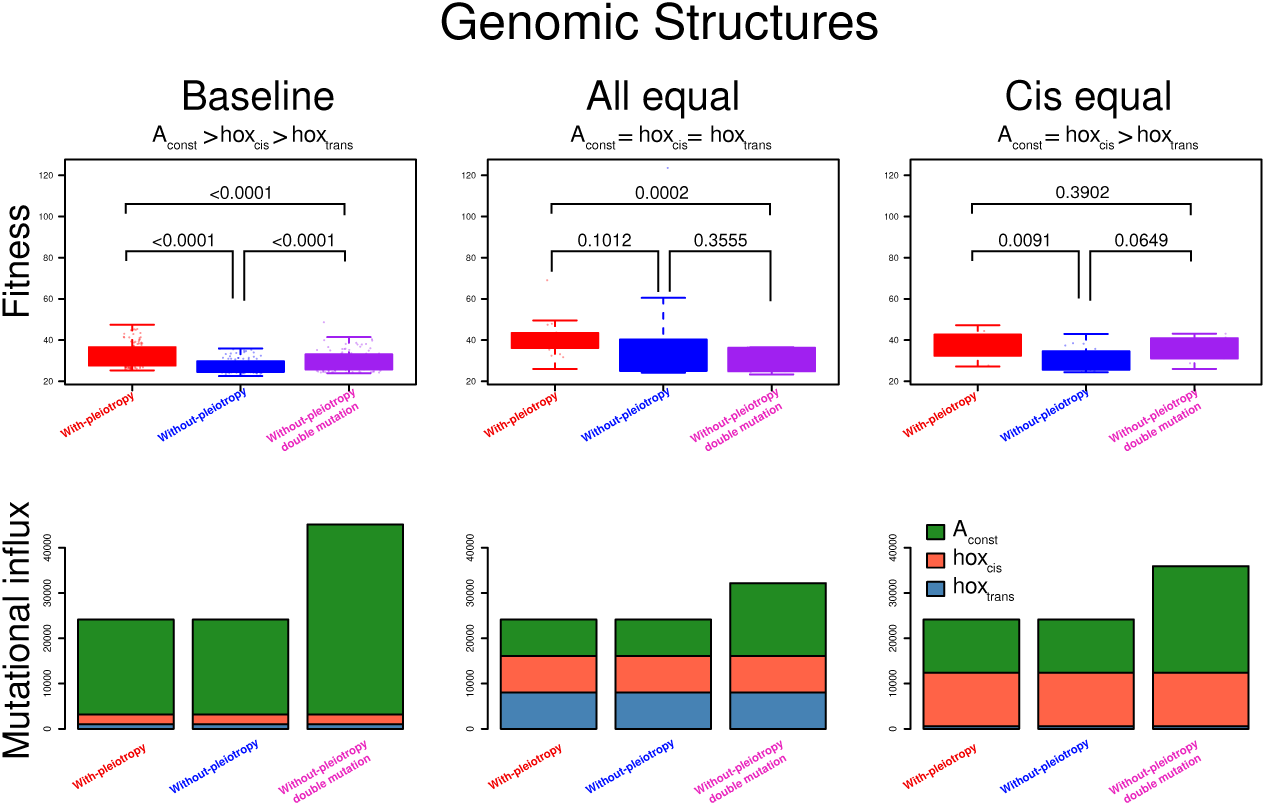
Genome structure. We tested two additional genomic structures, in which the relative mutational influx of the genetic elements differs from the baseline. Left: the baseline structure used in the main figures. Middle (“all equal”): all genetic elements have the same mutational influx. Right (“cis equal”): the **hox_cis_** elements have the same influx as the **A_constitutive_** elements, while the *hox_trans_*elements keep a lower influx. The bottom row shows the mutational influx in each case. In all structures tested, the with-pleiotropy setting reaches higher fitness than both without-pleiotropy settings, although not always significantly (Wilcoxon rank-sum tests; *p*-values are shown in the figure). Following the biological logic of our model, the “cis equal” structure corresponds to a situation in which Hox cis-regulatory elements make up a larger fraction of the genome, for example because more genes are under Hox control or their cis-regulatory regions are more extensive. The Hox gene itself still represents a small portion of the genome, so its mutation rate is kept low, as in the baseline. The “all equal” structure is biologically less plausible under this logic, since mutations in *hox_trans_* are expected to be strongly disruptive, as they affect several developmental parameters at once. This likely explains the lower fitness of the without-pleiotropy-double-mutation setting in this structure.

**Supplementary figure 5.**
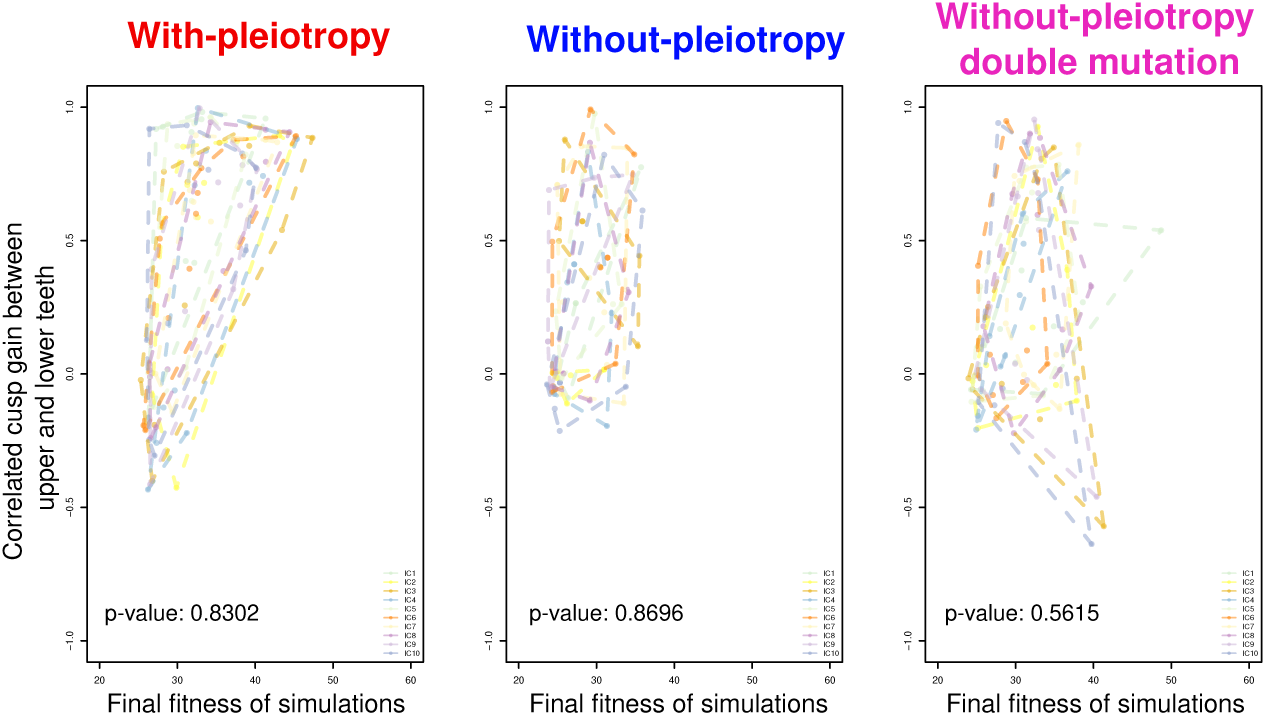
Relationship between correlated change of upper and lower teeth and the fitness achieved. Each setting is plotted separately, with simulations grouped by initial condition, as in Fig 3A. In none of the settings did the initial conditions significantly affect the relationship between correlated cusp change and final fitness. *p*-values are from *F* -tests comparing a linear model of final fitness on the correlation of cusp change with a model that also includes the initial condition.

**Supplementary figure 6.**
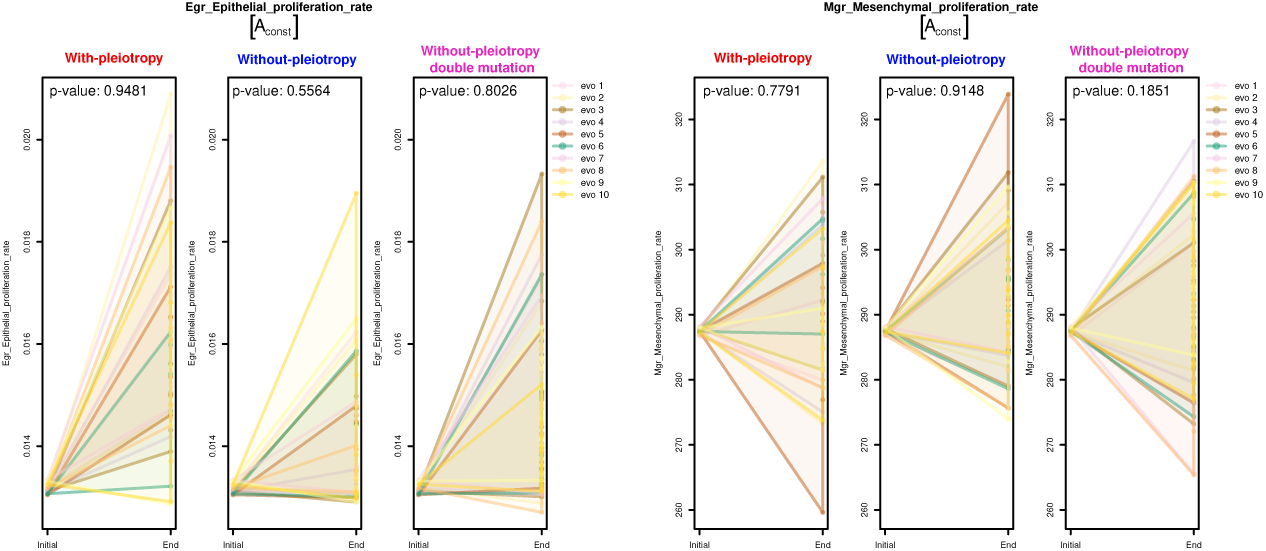
Final parameter values are largely independent of initial values. Raw values of two model parameters, epithelial proliferation and mesenchymal proliferation; shades indicate the different initial conditions. In each plot, the left side shows the mean parameter value at the start of the simulations of each initial condition, and the right side shows the mean final value of each simulation. Pearson correlations between initial and final values are shown in the plots; for these two parameters, they are not significant. Across all parameters (S1 Table5), only two of the 41 correlations in the with-pleiotropy setting remain significant after Benjamini–Hochberg correction (anterior bias and border growth, both **hox_cis_**), and none in the two without-pleiotropy settings.

**Supplementary figure 7.**
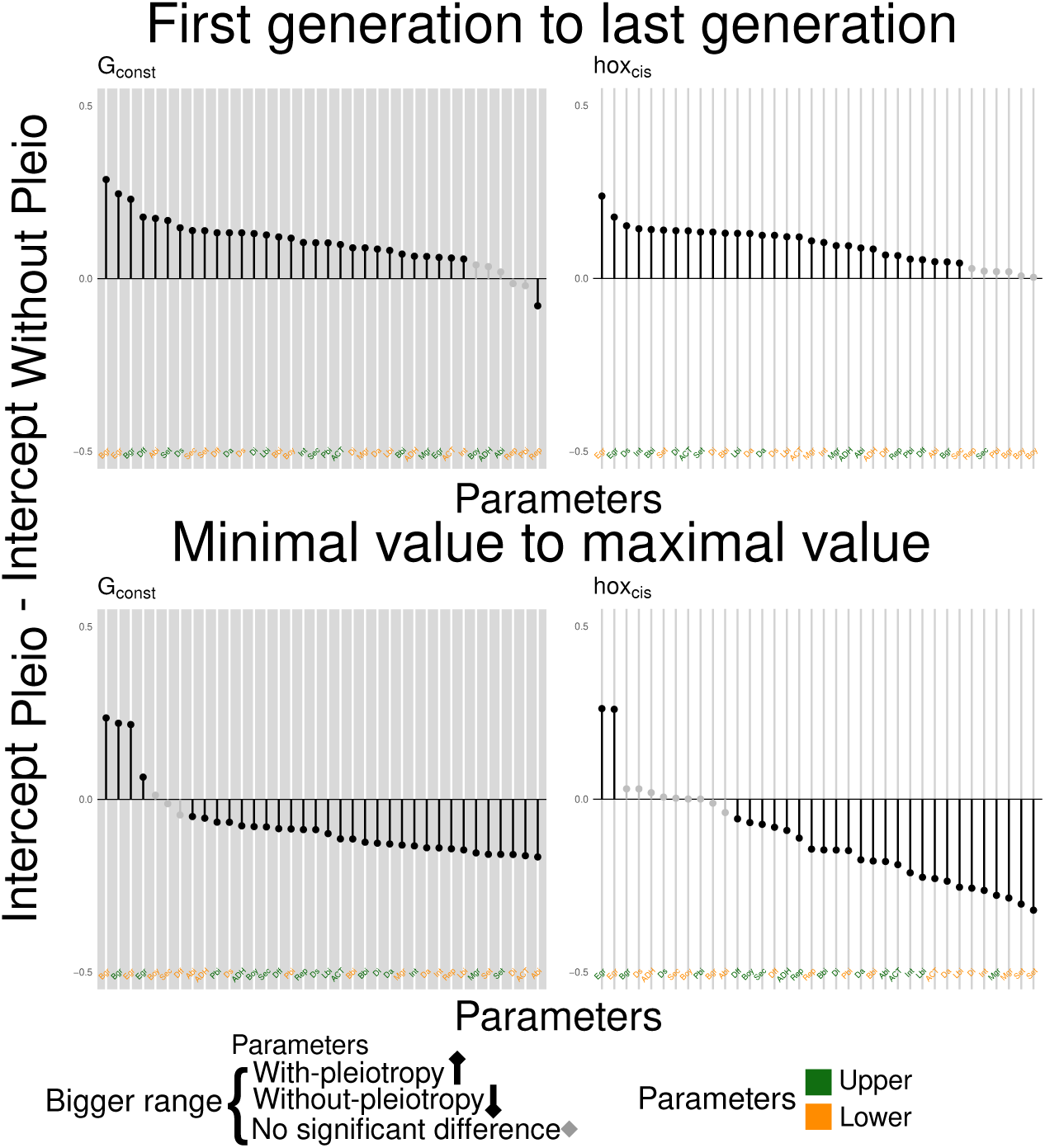
Intercept differences for each parameter. For each parameter, the difference in intercepts between the linear models of the with- and without-pleiotropy settings estimates the difference in parameter range at equal fitness (see Fig 5 and Methods, Ranges of genetic elements parameter change during evolution). Top: genetic divergence (Δ**_begin_***_−_***_end_**); bottom: overall genetic exploration (Δ**_min_***_−_***_max_**). The difference is calculated as the intercept with pleiotropy minus the intercept without pleiotropy, so positive values indicate a larger range in the with-pleiotropy setting. Gray bars indicate no significant difference between settings (Tukey’s multiple comparisons of the intercepts). Green and orange indicate the upper and lower teeth, respectively.

**Supplementary figure 8.**
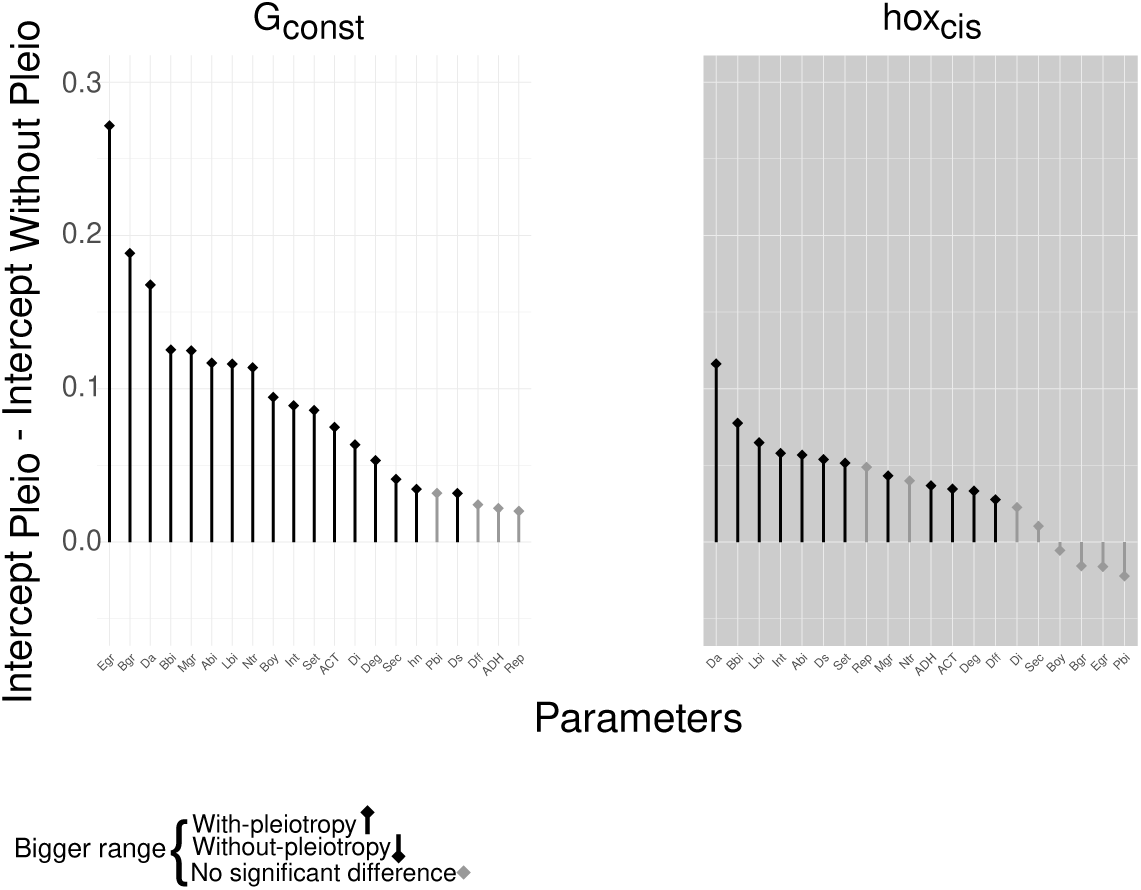
Adaptive changes for each parameter. For each parameter, we estimated the proportion of 50-generation windows in which the parameter’s evolution correlates with fitness improvement (adaptive windows), using a binomial generalized linear model with final fitness as covariate (see Methods, Adaptive change during evolution). Left: **A_constitutive_** elements; right: **hox_cis_** elements. Bars show the difference in predicted proportion of adaptive windows at equal fitness, calculated as the value with pleiotropy minus the value without pleiotropy; positive values therefore indicate more adaptive changes in the with-pleiotropy setting. Gray bars indicate no significant difference between settings (Tukey’s multiple comparisons in a linear model with final fitness as covariate). Adaptive changes are more frequent with pleiotropy for most parameters, in both elements.

**Supplementary figure 9.**
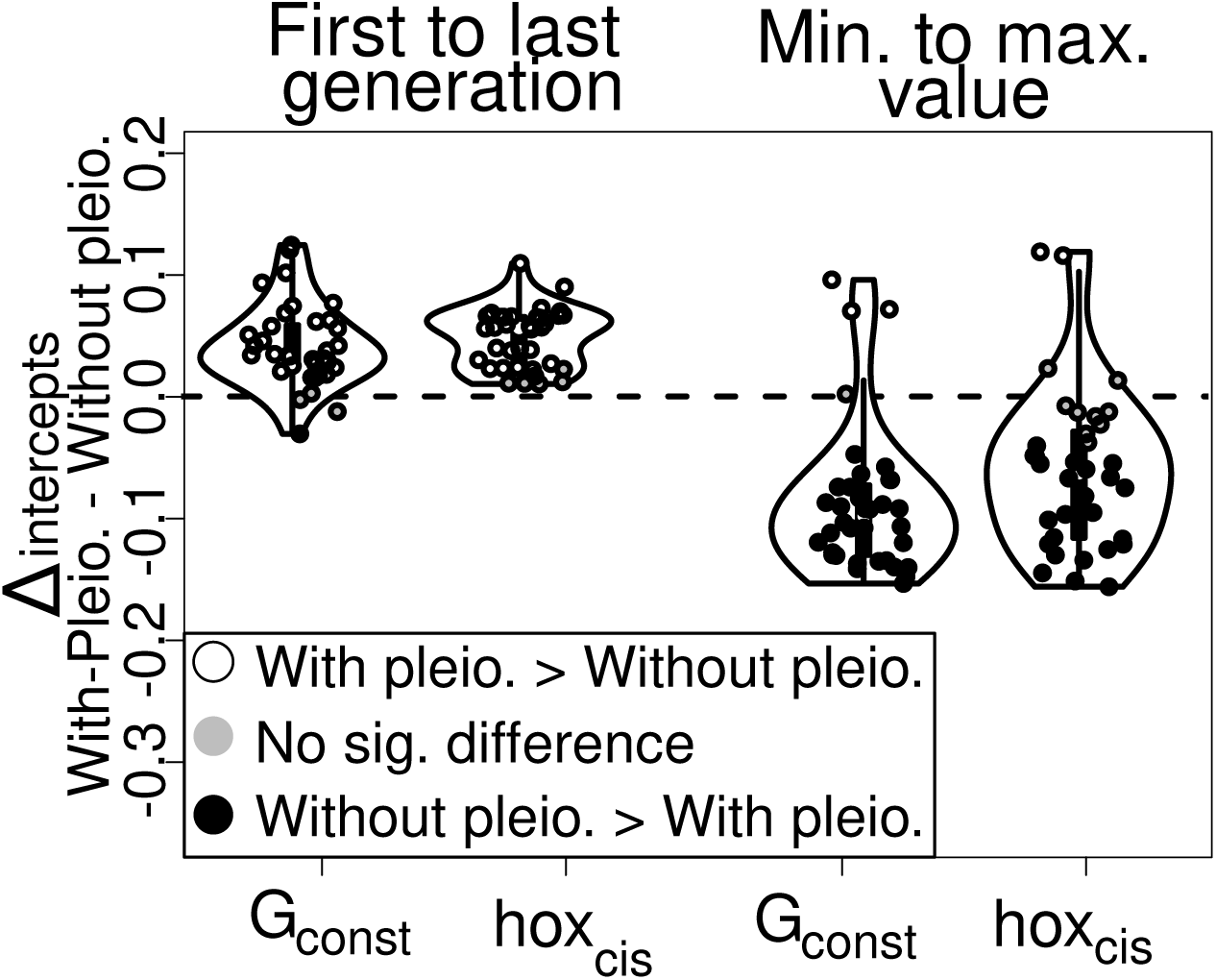
Intercept differences between the with-pleiotropy and without-pleiotropy-double-mutation settings. Same analysis as Fig 5B, comparing the with-pleiotropy setting with the without-pleiotropy-double-mutation setting. Each point is the difference between the intercepts of the linear models of the two settings (Δ**_intercepts_**), which estimates the difference in parameter range at equal fitness (see Methods, Ranges of genetic elements parameter change during evolution). White circles indicate a significantly larger range in the with-pleiotropy setting, black circles a significantly larger range in the without-pleiotropy-double-mutation setting, and gray circles no significant difference (Tukey’s multiple comparisons of the intercepts).

**Supplementary figure 10.**
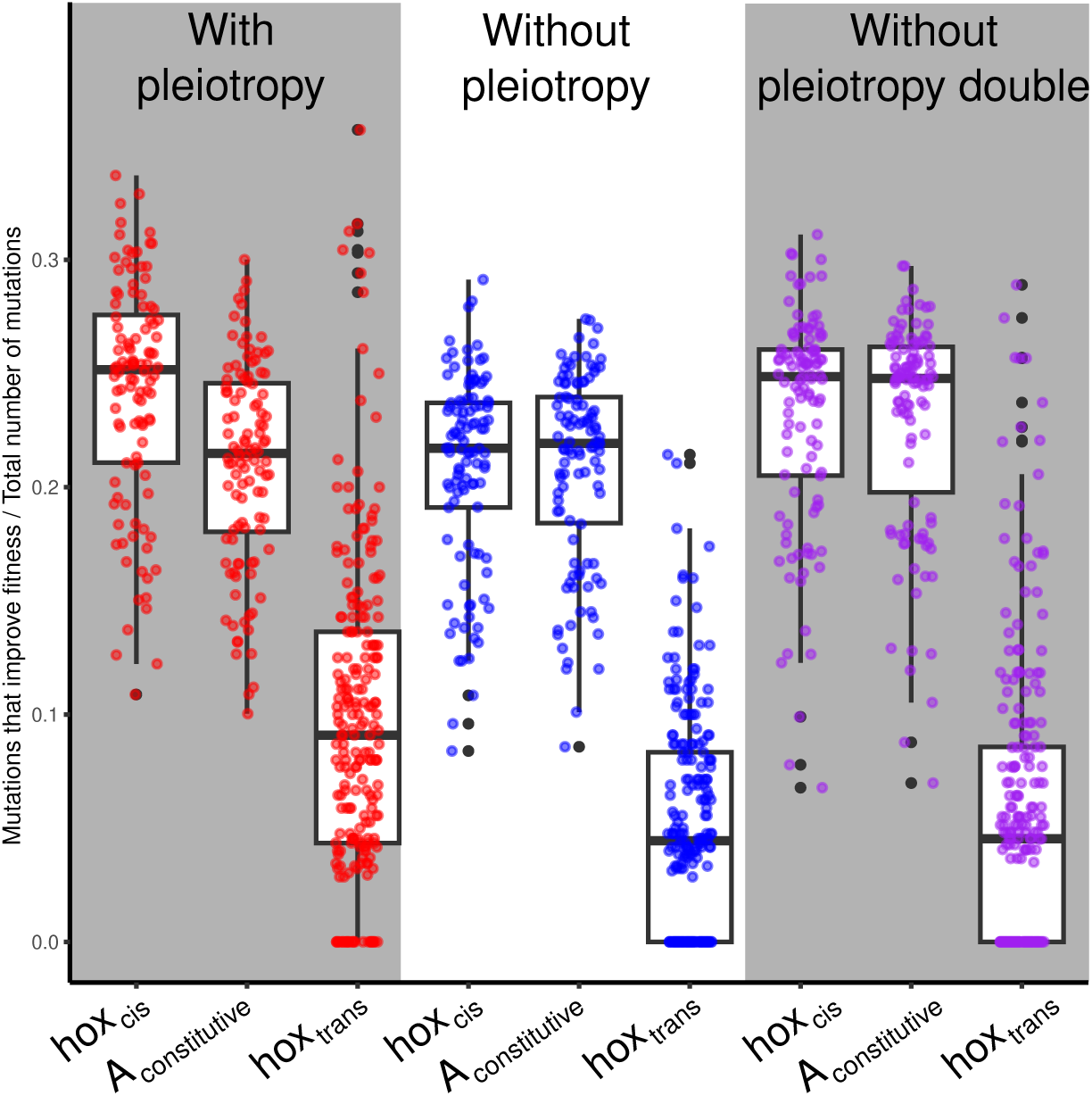
In the with-pleiotropy setting, mutations in the hox_cis_ parameters are more often adaptive. Same analysis as Fig 6, including the without-pleiotropy-double-mutation setting. The y-axis shows, for each simulation, the proportion of mutations that improved fitness and were passed on to the next generation. Mutations that disrupted development (i.e., for which the model failed to produce a final phenotype) are included in the total number of mutations. For simplicity, we only considered mutation events that changed one parameter at a time and combined the results for upper and lower teeth. In the without-pleiotropy-double-mutation setting, proportions are somewhat higher for both **hox_cis_** and **A_constitutive_**. Because mutations are more frequent in this setting, a larger fraction of mutation events affected more than one parameter and had to be discarded. These proportions are therefore based on a smaller subset of events and are not fully comparable with the other two settings.

**Supplementary figure 11.**
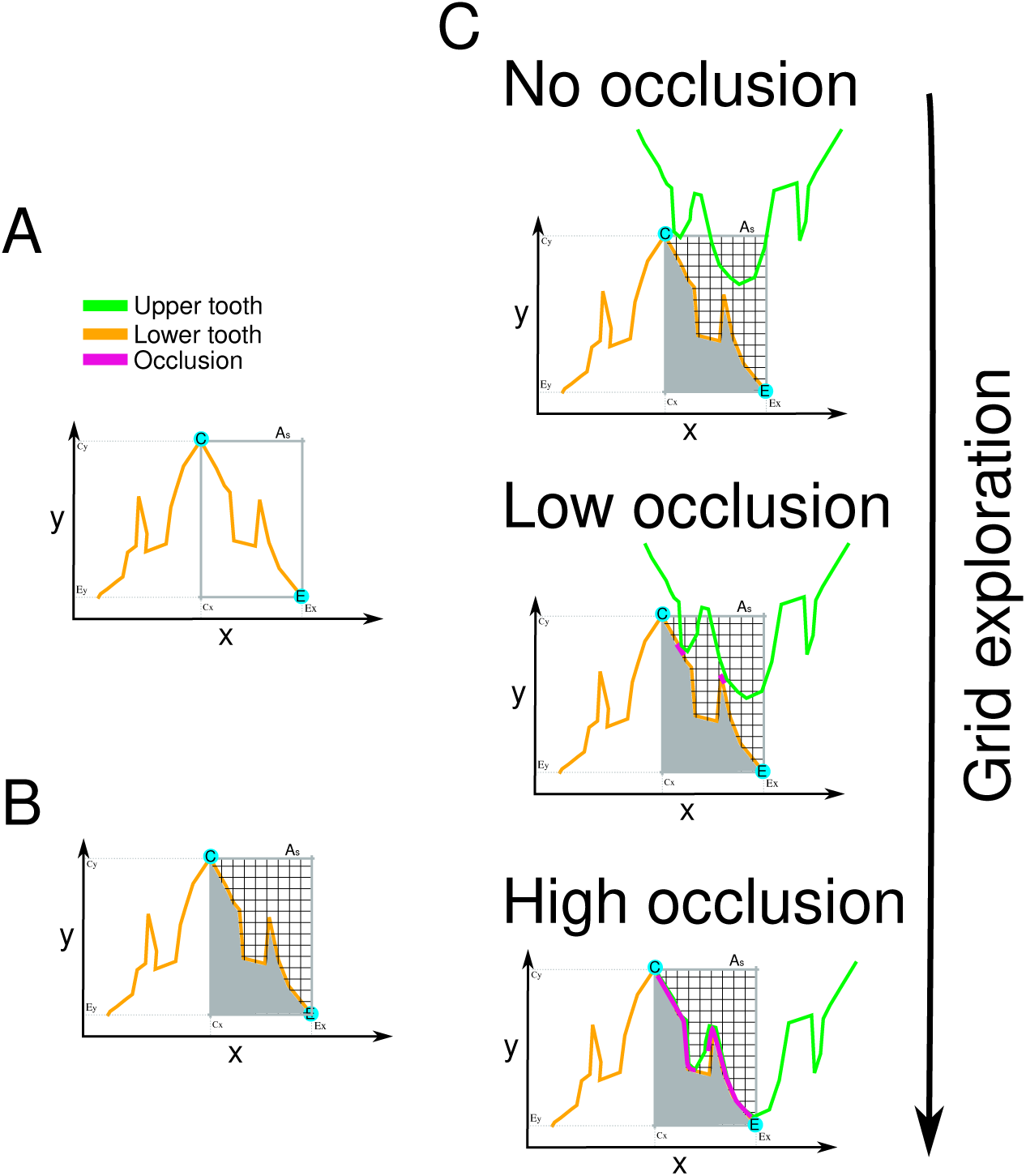
Occlusion measurement. A. To calculate occlusion, we first define a rectangular search area within which we find the position of the upper tooth that yields the largest occlusion. The search area *A_s_* is defined by two points of the lower tooth profile: its highest point *C* (the tip of the highest cusp) and the point with the largest *x*-value, *E*. B. The bottom and top edges of *A_s_* are set by the *y*-coordinates of *E* and *C* (*E_y_* and *C_y_*), and its right and left edges by their *x*-coordinates (*E_x_* and *C_x_*). Within *A_s_*, we exclude the area occupied by the lower tooth (gray) to prevent the two teeth from overlapping, a not very healthy arrangement. Intersections between the upper and lower tooth profiles are detected with the algorithm of Zerzan [74]. The remaining area is divided into a grid with a spacing of 1 model unit. C. The upper tooth profile is then translated so that its highest cusp is placed at the centre of each grid square in turn, with the rest of the profile following. For each translation in which no node of the upper profile falls within the excluded area, we calculate the contact between the two profiles. Points of the two profiles closer than 0.5 model units are considered to be in contact. The occlusion value is the sum, over both profiles, of the distances between consecutive contact points, i.e., the total length of the contact zone. The maximum value across all translations is the occlusion of the individual.

**S1 Table. Effect of initial conditions within each setting.** Kruskal–Wallis tests of cusp number (upper plus lower teeth), final fitness, cusp-change correlation and total acceleration as a function of initial condition (10 initial conditions, df = 9), for each setting at the baseline mutation rates, using one value per simulation. The table also gives *F* -tests comparing a linear model of final fitness on the cusp-change correlation with a model that also includes the initial condition (SFig 5). None of the tests is significant (all *p − values >* 0.05).

**S2 Table. Fitness-adjusted differences in parameter ranges between settings.** For each parameter, genetic element (**A_constitutive_**, **hox_cis_** and *hox_trans_*) and tooth, the table gives the difference in parameter range between settings at equal fitness (Δ**_intercepts_**), estimated from a linear model of range on final fitness and setting, for the genetic divergence (Δ**_begin_***_−_***_end_**) and the overall genetic exploration (Δ**_min_***_−_***_max_**). Differences are given for the with-pleiotropy setting minus each without-pleiotropy setting; positive values indicate a larger range with pleiotropy. *p*-values are from Tukey’s all-pairwise comparisons between the three settings (see Methods, Ranges of genetic elements parameter change during evolution). These values underlie Fig 5, SFig 7 and SFig 9.

**S3 Table. Tests of statistical assumptions.** Shapiro–Wilk tests of normality (per setting) and Levene’s tests of homoscedasticity (across settings) at the baseline mutation rates for cusp number (upper plus lower teeth), final fitness, upper- and lower-tooth EMD, cusp-change correlation and total acceleration, and for final fitness at the high and low mutation rates (SFig 3) and in the two alternative genome structures (SFig 4).

**S4 Table. Statistical results for the main and supplementary figures.** Fig 2: Wilcoxon rank-sum tests between settings for occlusion, upper- and lower-tooth EMD, and cusp number (upper plus lower teeth), computed on the ten best individuals of each simulation, with Benjamini–Hochberg-adjusted *p*-values. Fig 3: Spearman correlations between final fitness and the correlation of cusp change between upper and lower teeth, for each setting. Fig 4: *F* -tests comparing a linear model of total acceleration on final fitness with a model that also includes setting, and with a model that also includes the fitness *×* setting interaction, together with the coefficients of the additive model. Fig 6 and SFig 10: *t*-tests and Wilcoxon tests of the proportion of adaptive single mutations, paired within a setting and unpaired between settings. SFigs 3 and 4: Kruskal–Wallis and Benjamini–Hochberg-adjusted Wilcoxon rank-sum tests of final fitness between settings for each mutational regime.

**S5 Table. Correlation between initial and final parameter values.** Pearson correlations, per setting and parameter, between the initial value of each initial condition and the final value of each simulation, with Benjamini–Hochberg-adjusted *p*-values (correction across parameters within each setting; SFig 6).

## Notes

https://github.com/hagolani/PleiotropyDivergenceTwoOrgans

https://doi.org/10.5281/zenodo.23954307

### Competing Interest Statement

The authors have declared no competing interest.

### Summary of Updates

Improve by adding additional experiments to test tested the sensitivity of our model to variations in mutation rate, relative to mutation rates between genetic elements and initial conditions, and these new experiments confirmed the robustness of our results

